# Single-cell analyses reveal aberrant pathways for megakaryocyte-biased hematopoiesis in myelofibrosis and identify mutant clone-specific targets

**DOI:** 10.1101/642819

**Authors:** Bethan Psaila, Guanlin Wang, Alba Rodriguez Meira, Elisabeth F. Heuston, Rong Li, Jennifer O’Sullivan, Nikolaos Sousos, Stacie Anderson, Yotis Senis, Olga K. Weinberg, Monica L. Calicchio, NIH Intramural Sequencing Center, Deena Iskander, Daniel Royston, Dragana Milojkovic, Irene Roberts, David M. Bodine, Supat Thongjuea, Adam J. Mead

## Abstract

Myelofibrosis is a severe myeloproliferative neoplasm characterised by increased numbers of abnormal bone marrow megakaryocytes that induce progressive fibrosis, destroying the hematopoietic microenvironment. To determine the cellular and molecular basis for aberrant megakaryopoiesis in myelofibrosis, we performed high-throughput single-cell transcriptome profiling of 50,538 hematopoietic stem/progenitor cells (HSPCs), single-cell proteomics, genomics and functional assays. We identified an aberrant pathway for direct megakaryocyte differentiation from the earliest stages of hematopoiesis in myelofibrosis and associated aberrant molecular signatures, including surface antigens selectively expressed by *JAK2*-mutant HSPCs. Myelofibrosis megakaryocyte progenitors were heterogeneous, with distinct expression of fibrosis and proliferation-associated genes and putative therapy targets. We validated the immunoglobulin receptor G6B as a promising *JAK2*-mutant clone-specific antigen warranting further development as an immunotherapy target. Our study paves the way for selective targeting of the myelofibrosis clone and more broadly illustrates the power of single-cell multi-omics to discover tumor-specific therapeutic targets and mediators of tissue fibrosis.

## Introduction

Advances in single cell technologies have recently provided new insights into the cellular and molecular diversity and pathological mechanisms underlying many diseases, including cancers, pre-malignant and non-malignant conditions (Baslan and Hicks, 2017; Owen et al., 2018; Parikh et al., 2019). Parallel interrogation of mutation status and transcriptome at a single-cell level provide unprecedented opportunity to identify cancer cell-specific targets (Giustacchini et al., 2017; Rodriguez-Meira et al., 2019). Single cell resolution also uniquely enables identification of rare cell types and analysis of combinatorial patterns of gene expression, both of which are necessary to reconstruct differentiation trajectories and to accurately define cellular heterogeneity between populations such as normal and malignant tissues, as well as to identify the mediators of interactions between different cell types. For example, pathological fibrosis underlies many prevalent diseases including cancer, where fibrosis is well recognised to be important for disease progression and metastasis (Chandler et al., 2019; Cox and Erler, 2014). It is broadly proposed that pro-fibrotic mediators secreted by cancer cells and infiltrating immune cells activate non-malignant stromal cells such as myofibroblasts to deposit collagen fibrosis (Cox and Erler, 2014). However, an understanding of the specific cellular populations that mediate fibrosis in a given disease model, their molecular features, and the cellular pathways through which they are generated is necessary for these cells to be therapeutically targeted.

Myelofibrosis is a type of myeloproliferative neoplasms (MPNs) that results from somatic mutations in hematopoietic stem/progenitor cells (HSPCs) affecting MPL-JAK-STAT signaling, most commonly *JAK2V617F* (Kralovics et al., 2005; Levine et al., 2005). Myelofibrosis is characterised by progressive bone marrow fibrosis which destroys the hematopoietic microenvironment, resulting in the cardinal disease features of cytopenias, mobilization of HSPCs to peripheral blood, extramedullary hematopoiesis, and a high propensity for leukemia. Survival is typically 5-10 years from diagnosis and is not substantially improved by currently available drug therapies (O’Sullivan and Harrison, 2018). Megakaryocytes, the platelet-producing cells in the bone marrow, are dramatically increased in number in myelofibrosis and are the key cellular drivers of the destructive bone marrow remodelling via excessive release of pro-fibrotic cytokines and growth factors (Ciurea et al., 2007; Eliades et al., 2011; Martyre et al., 1997; Wen et al., 2015). In normal hematopoiesis, megakaryocyte progenitors (MkP) have a low proliferation rate, typically undergoing less than 8 cell divisions before mitotic arrest and the onset of polyploidization (Paulus et al., 2004). However, the cellular and molecular pathways giving rise to the dramatically increased megakaryocyte numbers and megakaryocyte dysfunction leading to tissue fibrosis in myelofibrosis are unclear.

In traditional models of hematopoiesis, megakaryocytes are said to arise from a bipotent progenitor shared with the erythroid (red cell) lineage, the megakaryocyte-erythroid progenitor (MEP) (Akashi et al., 2000; Debili et al., 1996; Kondo et al., 1997; Manz et al., 2002; Pang et al., 2005; Psaila et al., 2016; Psaila and Mead, 2019; Sanada et al., 2016). Recent advances in single-cell technologies including single-cell transplantation and lineage tracing studies of unperturbed hematopoiesis have revealed that hematopoiesis occurs over a continuum rather than via distinct, oligopotent intermediate steps (Laurenti and Gottgens, 2018; Psaila and Mead, 2019; Velten et al., 2017), and also that a proportion of HSCs, at least in the murine system, are megakaryocyte-biased but retain capacity for multilineage reconstitution (Adolfsson et al., 2005; Benz et al., 2012; Carrelha et al., 2018; Rodriguez-Fraticelli et al., 2018; Sanjuan-Pla et al., 2013; Shin et al., 2014). Lineage-committed megakaryocytes arising directly from HSCs, sometimes without cell division, have also been reported (Notta et al., 2016; Roch et al., 2015).

Targeting megakaryocytes in myelofibrosis has been shown to ameliorate the disease in mouse models and early-phase human studies (Eliades et al., 2011; Wen et al., 2015), but technical challenges have precluded extensive study of the cellular/molecular pathways for megakaryopoiesis in myelofibrosis. These include the rarity of megakaryocytes in the bone marrow, gaps in our knowledge of the cellular pathways of megakaryopoiesis and their extreme cell size and fragility. In addition, the severe fibrosis typically prevents bone marrow aspiration (“dry tap” aspirate). However, bone marrow HSPCs are mobilized to the peripheral blood in myelofibrosis. In this study, we utilized this phenomenon to capture peripheral blood HSPCs and perform the first in-depth study of abnormal megakaryocyte differentiation and function in myelofibrosis, suggesting novel cellular and molecular targets. Using multiparameter immunophenotyping, functional studies, high-throughput single cell transcriptome profiling (scRNAseq), targeted single cell mutational analysis with simultaneous scRNAseq (TARGET-Seq (Rodriguez-Meira et al., 2019)) and single cell proteomics we identify new potential targets for the inhibition of pathological megakaryocyte differentiation and megakaryocyte-induced fibrosis and validate G6B as a cell surface marker that may enable specific ablation of myelofibrosis cells using immunotherapy. This study illustrates the power of single cell ‘multi-omics’ in the characterisation of cellular heterogeneity in cancers associated with aberrant fibrosis, including the identification of novel therapeutic pathways and cancer cell-specific targets.

## Results

### Analysis of mobilized HSPCs demonstrates megakaryocyte-biased HSCs in myelofibrosis

Multiparameter flow cytometric analysis of the CD34^+^ lineage (lin)^−^ HSPC compartment in peripheral blood samples from healthy mobilized apheresis donors and patients with myelofibrosis (Suppl. Table 1) was performed to compare frequencies of the classically-defined HSPC subsets (Figure 1A). This demonstrated reduced lymphoid-primed multipotent progenitors (LMPP) and increased multipotent progenitors (MPP, Fig. 1A). The cell surface antigen CD41 has previously been reported to identify cells primed for megakaryocyte differentiation (Gekas and Graf, 2013; Haas et al., 2015; Psaila et al., 2016; Yamamoto et al., 2013). A 5-fold increase in the percentage of CD41^+^ cells was detected within both CD38-negative, early stem/progenitors (hematopoietic stem cell [HSC]/multipotent progenitor [MPP]) and CD38-positive, downstream progenitor (MEP/common myeloid progenitor [CMP]) cell fractions (Fig 1A, 1B), suggesting a bias towards megakaryocyte differentiation originating during the earliest phases of HSC lineage commitment. Morphological analysis of CD38^−^CD41^+^ and CD38^+^CD41^+^ cells from the CD34^+^lin^−^CD45RA^−^ compartment was consistent with undifferentiated blast cell morphology and not mature megakaryocytes (Suppl. Figs. S1A, S1B).

**Figure 1.**
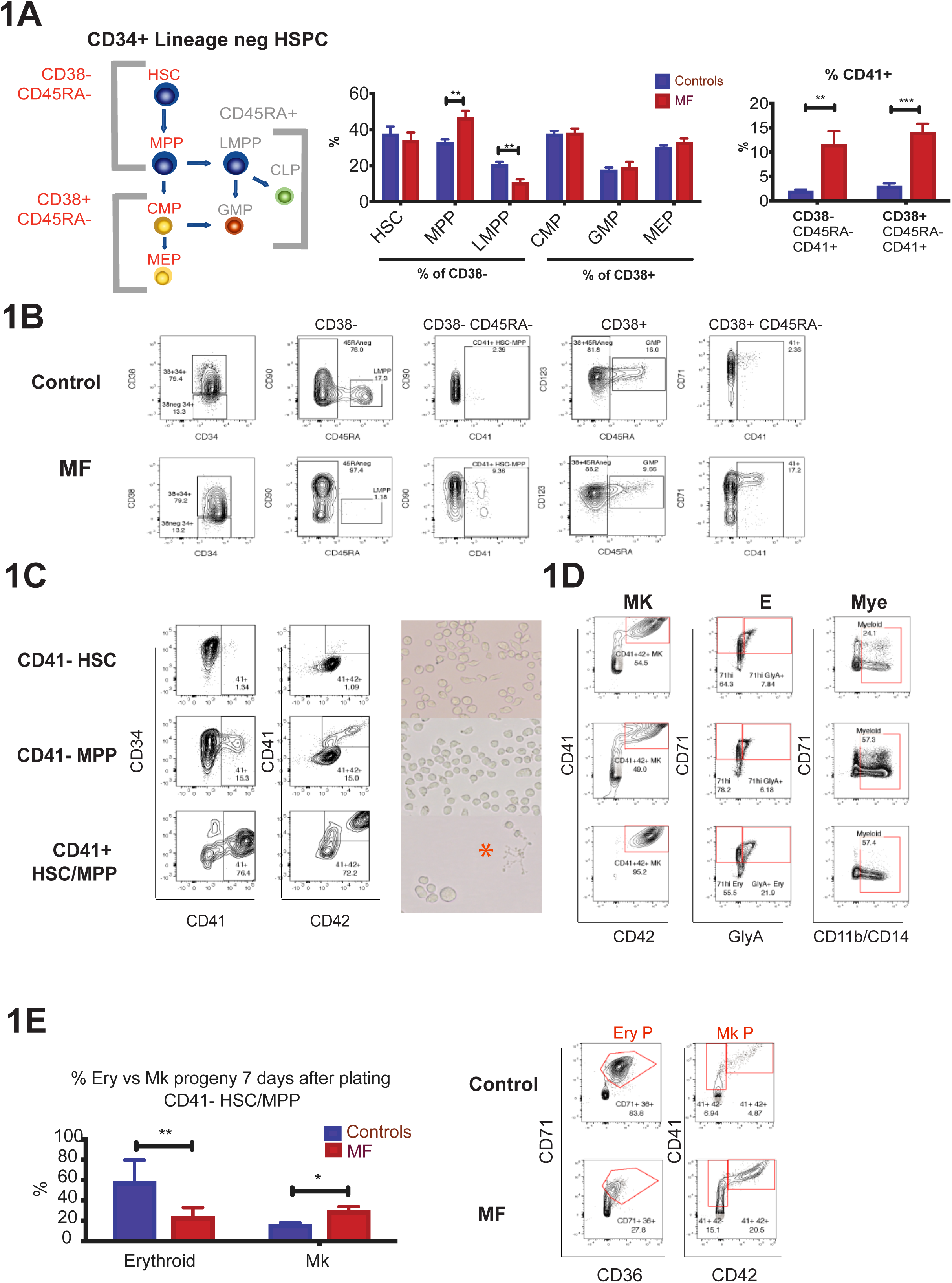
Multipotent myelofibrosis hematopoietic stem/progenitor cells (HSPCs) are biased for megakaryocyte differentiation. **1A: Left**: Model of classically-defined CD34^+^ lin^−^ HSPC subpopulations, in which multipotent stem cells (HSC – hematopoietic stem cells; MPP – multipotent progenitor cells) are CD38^−^ CD45RA^−^ and downstream progenitors (CMP - common myeloid progenitors; MEP – megakaryocyte-erythroid progenitors) are CD38^+^ CD45RA^−^. CD45RA^+^ populations (LMPP – lymphoid-primed multipotent progenitors; CLP – common lymphoid progenitors and GMP – granulocyte-monocyte progenitors) do not have erythroid or megakaryocyte potential. **Middle**: % of each classically-defined HSPC population in CD34^+^ lin^−^ compartment according to CD38 expression, demonstrating increased MPP and reduced LMPP in myelofibrosis. **Right**: % cells expressing CD41, a megakaryocyte surface antigen previously shown to identify cells with increased potential for megakaryocyte differentiation, is increased in myelofibrosis both in CD38^−^ CD45RA^−^ (HSC/MPP) and CD38^+^ CD45RA^−^ (CMP/MEP) compartments. (Myelofibrosis [MF] patients n=21; healthy donors n=17, see Supplementary Table 1.) **1B:** Representative FACS plot of a healthy donor control and myelofibrosis patient showing gating strategies. **1C: Left:** FACS analysis of CD41^−^ HSC (top), CD41^−^ MPP (middle) and CD41^+^ HSC/MPP (lower) from healthy donors cultured in megakaryocyte differentiation media (+rhTPO & SCF). CD41^+^ HSC/MPP had increased potential for megakaryocyte differentiation, with faster acquisition of the mature megakaryocyte antigen CD42 at an early timepoint (day 6). **Right:** images of cultures showing enlarged cell size and evidence of proplatelet formation (red star) indicative of accelerated megakaryocyte differentiation from CD41^+^ HSC/MPP. Representative examples of 3 replicate experiments shown. **1D**: FACS analysis of CD41^−^ HSC (top), CD41^−^ MPP (middle), and CD41^+^ HSC/MPP (lower) from healthy donors cultured for 12-14 days in megakaryocyte (MK), erythroid (E) or myeloid (Mye) differentiation media. CD41^+^ HSC/MPP showed a higher % of mature CD41^+^42^+^ megakaryocytes and glycophorin A^+^ CD71^+^ erythroblasts, and equivalent CD11b/CD14^+^ myeloid cells *vs*. CD41^−^ HSC and MPP. Representative examples of 3 replicate experiments shown. Percentages shown are the % of total live, single cells analysed (7AAD^−^, doublets excluded). **1E:** CD41^−^ HSC/MPP cultured in ‘bipotent’ erythroid and megakaryocyte differentiation media showed a bias towards megakaryocyte (Mk P) *vs*. erythroid (Ery P) differentiation in myelofibrosis as compared to healthy donor control cells. Left – summary chart (n=3 for each of myelofibrosis and contols); right – example FACS plots. (Chart shows mean+SEM.**P=0.01; *P=0.05, controls-n=3;myelofibrosis [M]),n=4). Charts show Mean+SEM,***P<u><0</u>.001; **P≤0.01; *P<0.05; for multiple t-test with FDR for Benjamini Hochberg correction where appropriate). See also Suppl. Fig. S1.

The CD41^+^ fraction of human CD38^+^ CD34^+^ lin^−^ CD45RA^−^ HSPCs contains megakaryocyte-biased progenitors with significant erythroid differentiation potential as well as unipotent MkP (Miyawaki et al., 2017; Psaila et al., 2016). However, the phenotype of CD41^+^ cells within the CD38^−^ HSC/MPP compartment has not previously been defined. We therefore first sought to determine whether the CD41^+^ HSC and MPP cells isolated from healthy donors retained capacity for multilineage differentiation or were lineage-committed MkP. CD34^+^ Lin^−^ CD38^−^ CD45RA^−^ CD90^+^ CD41^−^ (CD41-HSC), CD34^+^ Lin^−^ CD38^−^ CD45RA^−^ CD90^−^ CD41^−^ (CD41-MPP) and CD34^+^Lin^−^ CD38^−^ CD45RA^−^ CD41^+^ (CD41^+^HSC/MPP) cells were isolated by fluorescence-activated cell sorting (FACS, Suppl. Fig. S1A) for liquid culture differentiation assays. Stimulated with thrombopoietic cytokines, CD41^+^ HSC/MPP cells showed accelerated megakaryocyte differentiation with a substantially higher proportion of cells expressing the mature megakaryocyte surface antigen CD42, a large cell size and proplatelet extensions at early timepoints as compared to CD41^−^ HSC and MPP (Fig. 1C). In combined megakaryocyte, erythroid and myeloid differentiation assays, CD41^+^ HSC/MPP not only showed increased megakaryocyte differentiation but also a similar potential for CD11b/CD14^+^ myeloid differentiation and superior potential for CD71^+^/glycophorin A erythroid differentiation than CD41^−^ fractions (Fig. 1D).

In comparison to those from healthy donors, CD41^−^ HSC/MPP cells from myelofibrosis patients showed significant megakaryocyte *vs.* erythroid bias (Fig 1E), in keeping with the clinical phenotype of myelofibrosis patients in which excessive megakaryocyte numbers occur in parallel with anemia. In single-cell clonogenic assays supportive of myeloid and erythroid (but not megakaryocytic) colony formation (methocult), CD41^+^ and CD41^−^ fractions of HSC and MPP gave rise to expected colony frequencies with no significant difference between healthy donors and myelofibrosis patients (Suppl. Fig. S1C). Together, these results support that in myelofibrosis, HSPCs are biased towards megakaryocyte-lineage differentiation from the earliest stem cell compartment, before expression of canonical megakaryocytic markers.

### High-throughput single cell RNA-sequencing identifies a distinct pathway for megakaryocyte differentiation in myelofibrosis

To identify the cellular and molecular basis for megakaryocyte-biased hematopoiesis in myelofibrosis without bias from pre-selected cell surface antigens, high-throughput scRNAseq was performed on 48,421 individual CD34^+^ lin^−^ HSPCs from patients with JAK2V617F+ post-polycythaemia myelofibrosis (30,088 cells, n=3) according to WHO criteria (Arber et al., 2016) and age-matched healthy donors (18,333 cells, n=2) using the 10x Genomics Chromium platform (Suppl. Table 2). Following filtering, quality control and exclusion of doublets, 47,804 cells passed quality control (29,536 myelofibrosis and 18,249 control, Suppl. Table 3). Healthy donor control and myelofibrosis cells were aggregated separately and the donor effect was regressed out (Suppl. Fig. S1D).

Dimensionality reduction and unsupervised clustering were performed using a uniform manifold approximation and projection (UMAP) method combined with k-means clustering to enable identification of distinct cell populations while preserving inter-cluster relationships (Becht et al., 2018), Fig 2A). Clusters were identified by analysis of differentially expressed genes for each cluster (Fig. 2A, Suppl. Figs. S1E & 2, Suppl. Tables 4, 5). “Lineage signature gene sets” were then defined containing genes selectively associated with erythroid, myeloid, lymphoid and megakaryocyte lineages (Suppl. Fig. S2, Suppl. Table 6) and superimposed on the UMAPs (Fig. 2B). This highlighted two distinct clusters of cells expressing megakaryocyte signature genes among myelofibrosis CD34^+^lin^−^ HSPCs, accounting for around 15% of the HSPCs overall. In contrast, very few healthy donor control HSPCs expressed megakaryocyte lineage signature genes and did not form a distinct cluster but were scattered within the erythroid cluster (Fig. 2B (inset), Suppl. Fig. S1E).

**Figure 2:**
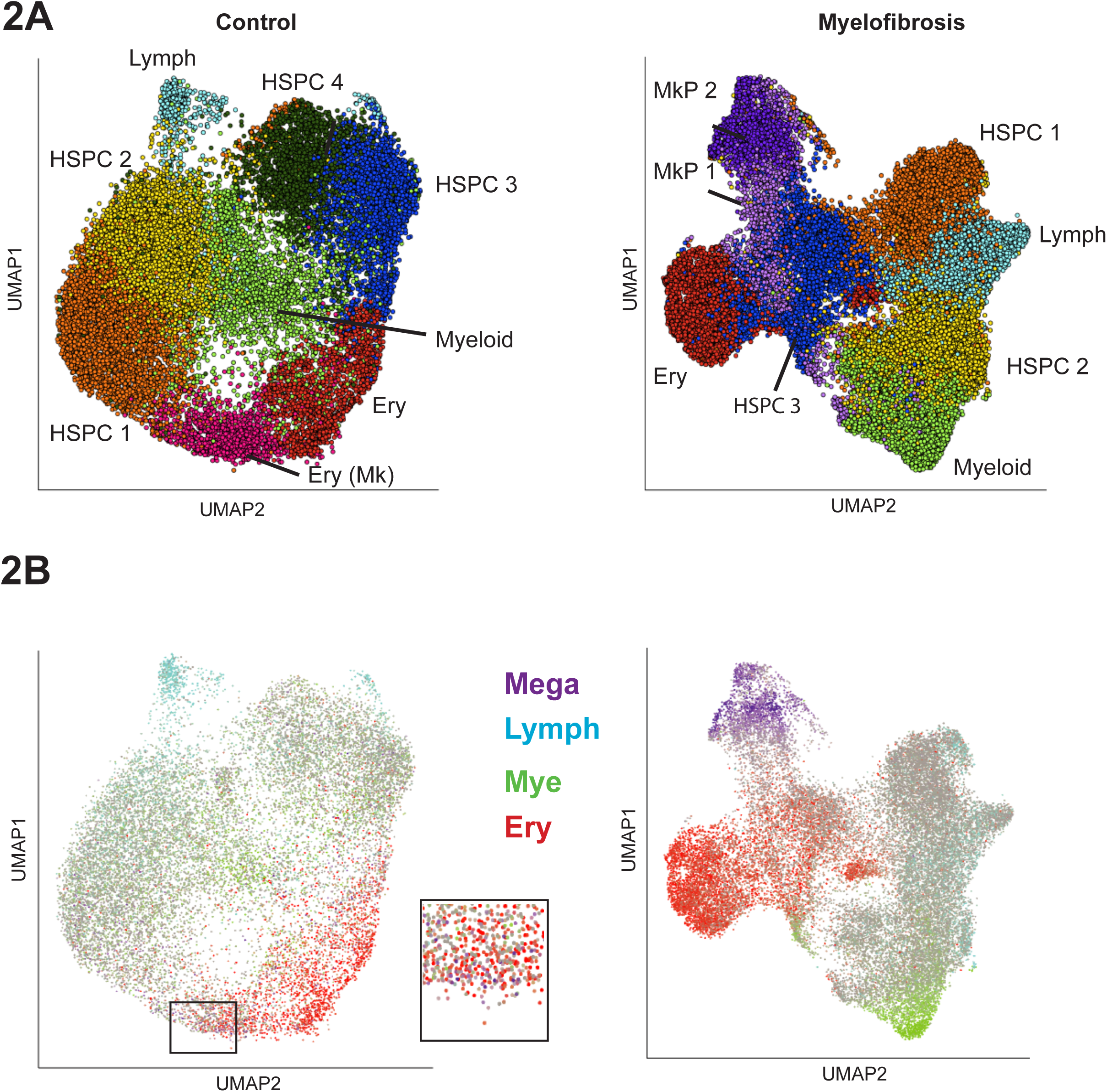
High-throughput single cell RNA-sequencing of 47,804 CD34+Lin-HSPCs reveals marked expansion of megakaryocyte-progenitors (MkP) in myelofibrosis. **2A:** Dimensionality reduction using Uniform Manifold Approximation and Projection (UMAP) on batch-corrected aggregates of control (n=18,249) and myelofibrosis (n=29,536) cells identified distinct cell populations while preserving inter-cluster relationships. Cells were partitioned using k-means clustering and annotated according to expression of key marker genes for the major cell hematopoietic cell types. Distinct HSPC subsets expressing genes associated with myeloid, lymphoid and erythroid lineages were identified within both healthy donor and myelofibrosis cell aggregates, while distinct MkP clusters were present only among myelofibrosis HSPCs. HSPC clusters showing no evidence of lineage priming are labelled HSPC 1-4. Patients with *JAK2V617F*+ myelofibrosis (n=3) and age-matched controls (n=2) were used (see also Suppl. Fig. S1D, S1E and Suppl. Table 2). **2B:** Expression of lineage signature gene sets for the 4 major hematopoietic lineages were superimposed on the UMAP (megakaryocyte-purple; lymphoid – blue; myeloid – green; erythroid – red; grey – uncommitted or expression of >1 lineage gene set). Inset shows higher magnification view of infrequent MkP cells present (purple cells) within an erythroid (red cells) cluster in the control UMAP in contrast to the distinct MkP cluster in the myelofibrosis aggregate. Abbreviations: HSPC – hematopoietic stem and progenitor cells; Mye – myeloid; Lymph-Mye – lymphoid/myeloid; Ery – erythroid; Mega – megakaryocyte. See also Suppl. Fig. S1E.

To study differentiation trajectories, cells were ordered in gene expression space using forced directed graphs and lineage signature gene scores superimposed on the graphs (Fig. 3A). Myeloid, erythroid and lymphoid trajectories were observed in both healthy donors and myelofibrosis patients. Expression of megakaryocyte genes (purple) was observed along a distinct trajectory arising directly from uncommitted HSPCs (grey) in addition to along the erythroid trajectory (red) only in myelofibrosis HSPCs (Fig. 3A & B, right). By contrast, in healthy donor controls, expression of megakaryocyte genes occurred only within the same trajectory as the erythroid genes (Fig. 3B, left). Aggregating all 47,804 control and myelofibrosis cells together demonstrated that 2568/2575 (99.7%) of cells within the megakaryocyte trajectory derived from myelofibrosis patients, with an almost complete absence of healthy donor cells (7/2575 cells, 0.3%, Fig. 3C). Together with functional data (Fig. 1), these data suggest a model in which a direct route for MkP production from HSPC is massively expanded in *JAK2V617F* mutation positive myelofibrosis, in addition to increased production of megakaryocytes via a shared trajectory with the erythroid lineage (Fig. 3B, Fig. 3D).

**Figure 3:**
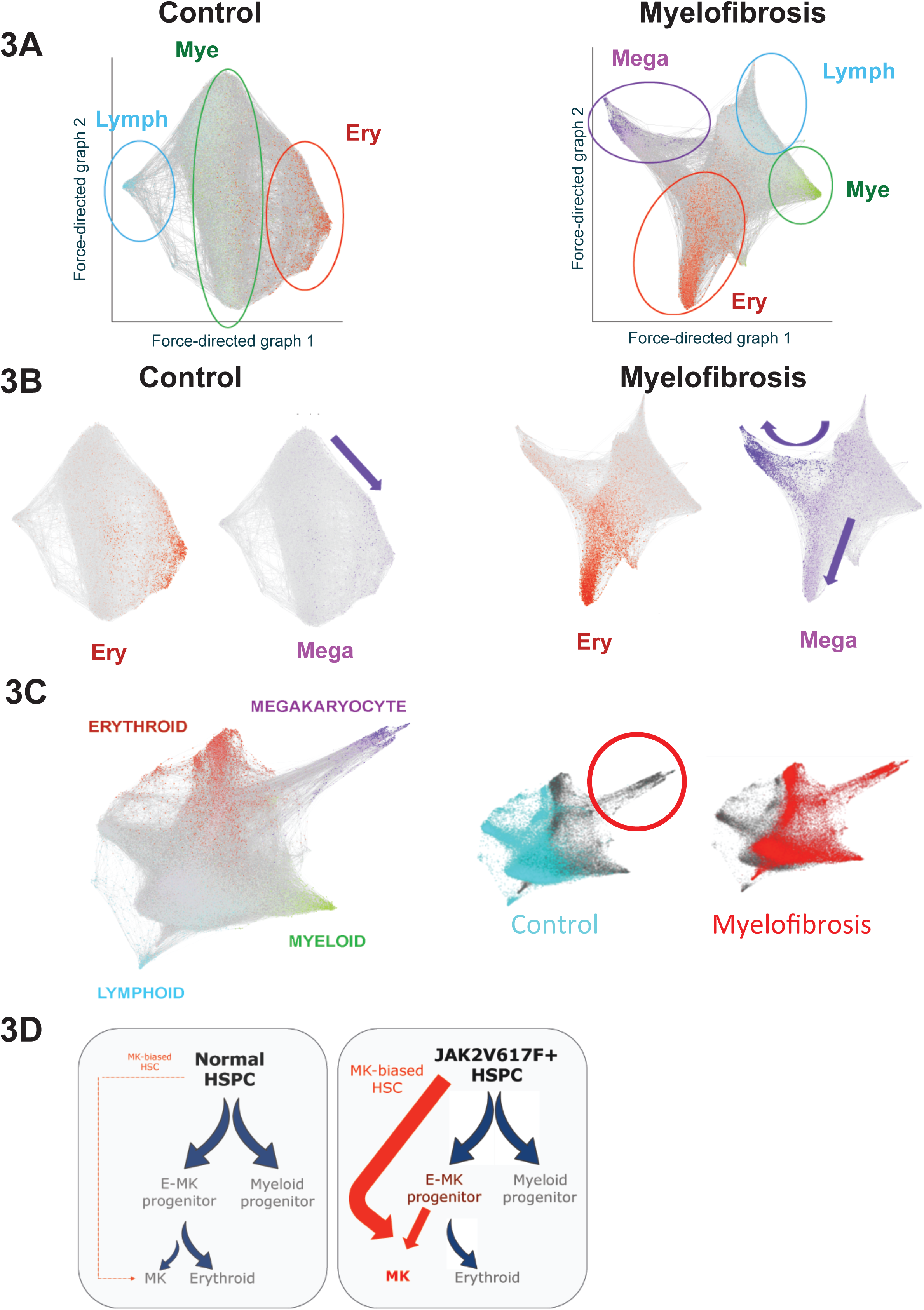
A distinct trajectory for direct emergence of megakaryocyte progenitors from uncommitted HSPCs in myelofibrosis. **3A:** Force-directed graphs for healthy donor control and myelofibrosis cell aggregates and gene expression trajectories visualized by superimposing the expression scores of myeloid (mye - green), erythroid (ery – red), lymphoid (lymph – blue) and megakaryocyte (mega – purple) lineage-signature gene sets. Grey cells respresent uncommitted HSPC or cells with expression of >1 lineage signature. A distinct megakaryocyte trajectory is evident only in the myelofibrosis graph. Each cell is represented by a node (dot) and the edges (lines) connect all pairs of cells that share at least five nearest neighbors. **3B:** Expression of megakaryocyte signature genes occurs along the same trajectory as erythroid genes in the control cell aggregate (left, purple arrow indicates megakaryocyte trajectory), whereas an expanded megakaryocyte differentiation path is evident both along a shared erythroid-megakaryocyte trajectory as well as in a distinct trajectory in the myelofibrosis aggregate (right, purple arrows indicate megakaryocyte trajectories). **3C:** Aggregating all 47,804 control and myelofibrosis cells together (**left**) and labelling the cells according to disease state (**right**) demonstrates almost complete absence of control cells in the direct megakaryocyte differentiation trajectory. **3D:** Proposed model for *JAK2V617F*-driven hematopoiesis in myelofibrosis, showing increased generation of megakaryocyte progenitors via the normal shared megakaryocyte-erythroid pathway as well as via an aberrant ‘direct’ route for megakaryopoiesis.

### Identifying molecular drivers for aberrant megakaryopoiesis in myelofibrosis

To identify potential molecular drivers for the aberrant megakaryocyte differentiation trajectory, we performed unsupervised k-means clustering on a 3-dimensional k-nearest neighbor (KNN) graph aggregate of all 47,804 cells (see .html file, Suppl. Item 1) and identified the paths taken by cells from the earliest undifferentiated HSPCs along the aberrant megakaryocyte trajectory that comprised almost entirely of myelofibrosis cells and the erythroid/megakaryocyte trajectory containing both myelofibrosis and control cells (Fig. 4A). Expression patterns of 1,639 human transcription factors (Lambert et al., 2018) were examined along the two trajectories and genes clustered according to patterns of change in gene expression levels. Transcription factor genes showing progressive changes, either increased or decreased expression, along the two trajectories were further inspected (Suppl. Figs. S3, S4) and compared between the two trajectories (Fig. 4B). Expected patterns of expression of transcription factors known to be involved in megakaryocyte and erythroid differentiation were observed (e.g. progressive increase in *GATA1, GATA2*), as well as antagonistic expression of two key regulators of megakaryocyte-erythroid cell fate decision *FLI1* and *KLF1* (Bouilloux et al., 2008; Dore and Crispino, 2011; Frontelo et al., 2007; Palii et al., 2019; Siripin et al., 2015) (Fig. 4B, 4C). Additional genes not previously implicated as regulators of megakaryocyte vs. erythroid differentiation showed striking differential expression between the trajectories, included *YBX1, PLEK, SOX4* and *MYC* (Fig. 4B, 4C), suggesting additional novel targets for strategies to specifically inhibit pathological megakaryopoiesis while preserving erythropoiesis in myelofibrosis patients.

**Figure 4:**
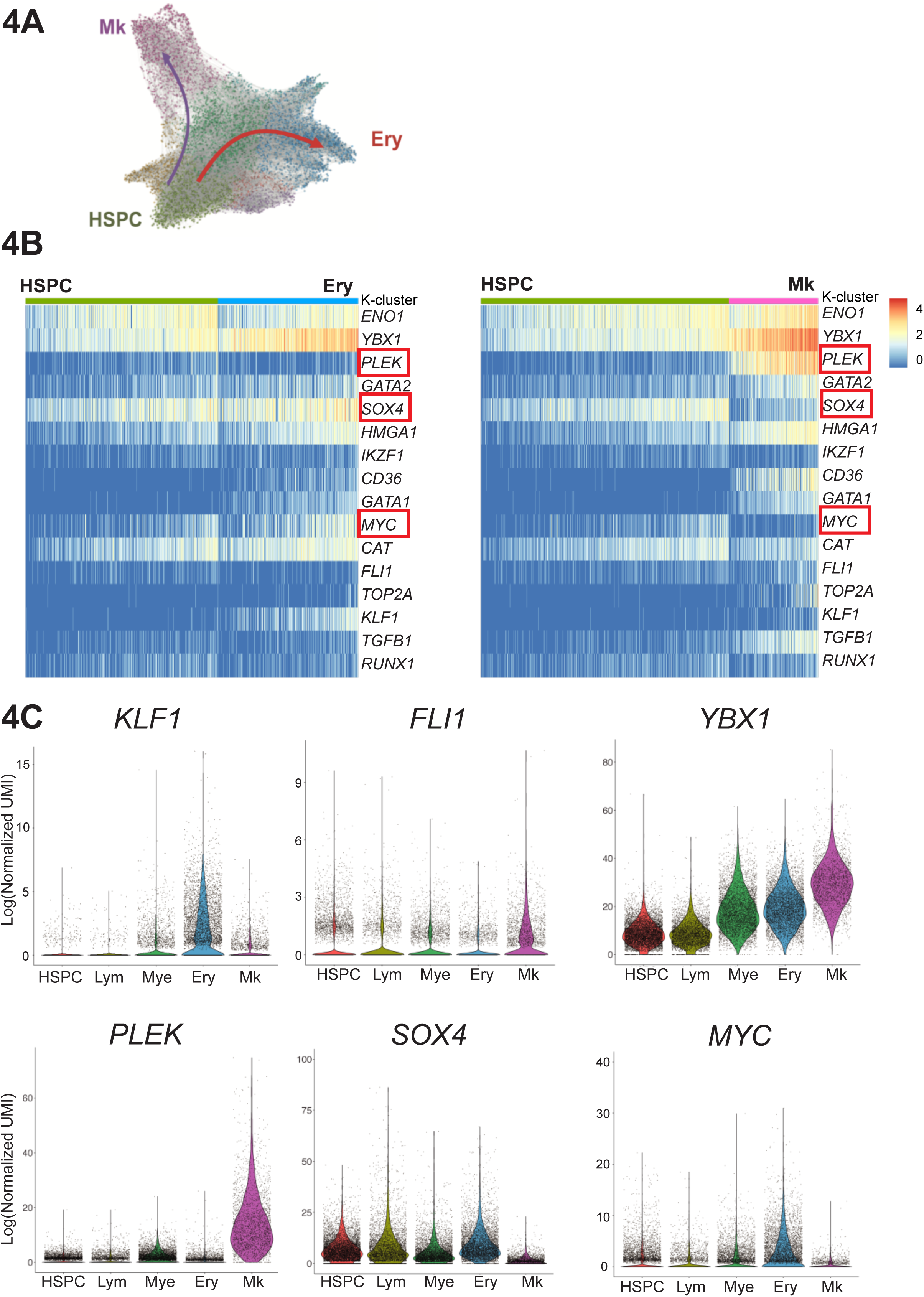
Unique molecular drivers of aberrant megakaryocyte differentiation. **4A:** Unsupervised k-means clustering on the k-nearest neighbor (KNN) graph aggregate of all 47,804 control and myelofibrosis cells. Differentially expressed genes for each cluster were used to identify erythroid (Ery, blue), megakaryocyte (Mk, purple) and uncommitted HSPC clusters (HSPC, green). See also 3D graph in .html file, Suppl. Item 1. **4B:** Expression of 1,639 known/putative human transcription factors were examined along the Ery pseudo-temporal path and the Mk path. Selected genes showing distinct expression along the aberrant HSC→ Mk (myelofibrosis only) and HSC → Ery (control and myelofibrosis) pseudo-temporal paths are shown in the heatmaps. *PLEK* is selectively upregulated and *SOX4* and *MYC* downregulated along HSC→ Mk, with substantially higher *YBX1* expression along the Mk trajectory than erythroid. See also Suppl. Figs. S3 and S4. **4C:** Violin plots for transcription factors *KLF1* and *FLI1*, known regulators of Mk/Ery cell fate specification as well as novel additional potential molecular regulators of Ery vs. Mk only differentiation (*PLEK, SOX4, MYC, YBX1)* are shown for clusters expressing genes associated with lymphoid (Lym), myeloid (Mye), erythroid (Ery) and megakaryocyte (Mk) lineage differentiation and the initial HSPC cluster on the KNN trajectory plot shown in 4A. 3 outlying cells not shown (MYC = 2 cells; SOX4 = 1 cell).

### Identifying mediators of megakaryocyte-induced fibrosis

To evaluate the pathological role of the expanded population of MkP in driving bone marrow fibrosis, we next examined potential mediators of fibrosis. Fibrosis regulators were identified from previously published datasets studying lung and liver fibrosis as well as bone marrow fibrosis (Allen et al., 2017; Blackman et al., 2013; Corvol et al., 2015; Gu et al., 2009; Mondet et al., 2015; Mushiroda et al., 2008; Noth et al., 2013; Ulveling et al., 2016; Wattacheril et al., 2017; Wright et al., 2011). Genes detected at expression levels over 1 (using log-transformed UMI) were selected for a ‘fibrosis signature’ gene score (Suppl. Table 6). Superimposition of this score on the UMAPs for healthy donor and myelofibrosis HSPCs clearly highlighted the myelofibrosis MkP cluster cells (Fig. 5A). All healthy donor and myelofibrosis cells expressing at least two megakaryocyte signature genes were then extracted for further analyses. Importantly, *TGFB1* was detected both in a higher fraction of myelofibrosis MkP than healthy donor MkP (58.6% *vs*. 36.5%) and also expressed at substantially higher levels per cell (Fig. 5B). This indicates that megakaryocyte-induced fibrosis in myelofibrosis is due to an aberrant pro-fibrotic megakaryocyte phenotype in addition to increased megakaryocyte numbers, an observation which would not have been possible without single-cell analysis. *LTBP1*, which encodes a protein that targets the latent form of transforming growth factor beta (TGFβ) and contributes to its activation (Robertson et al., 2015), showed a similar pattern with expression detected in a substantially higher % of myelofibrosis MkP as well as increased expression per cell (Fig. 5B).

**Figure 5:**
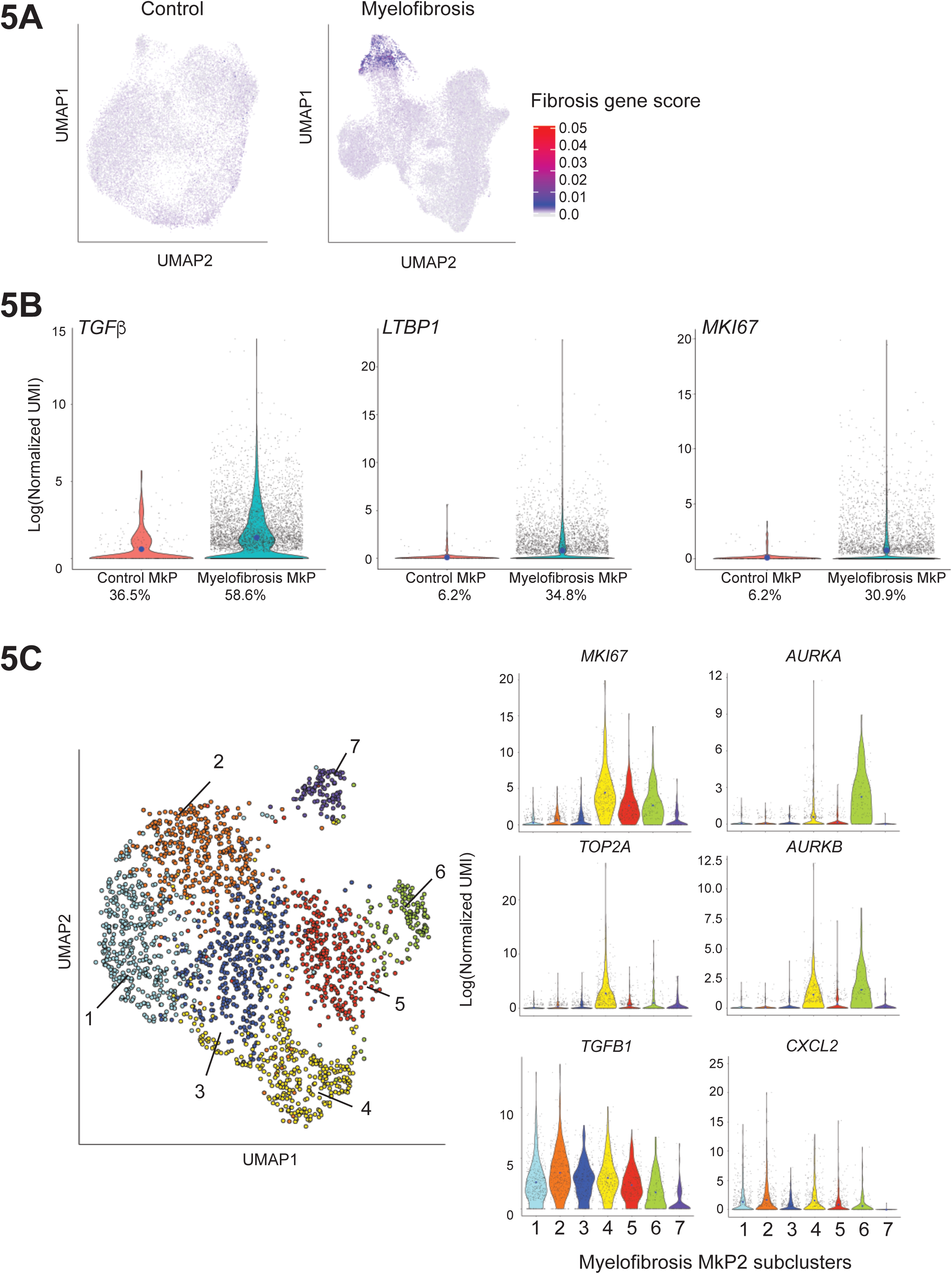
Myelofibrosis-specific megakaryocyte progenitors (MkP) strongly express mediators of tissue fibrosis. **5A:** Expression of a 14-gene ‘fibrosis score’ (see also Suppl. Table 6) derived from previously published datasets examining bone marrow, liver and lung fibrosis superimposed on UMAP plots of healthy donor control and myelofibrosis aggregates (from Figure 2B) identifies high level of expression of pro-fibrotic mediators by myelofibrosis MkP (MF-MkP). **5B**: MkP identified as HSPCs expressing ≥ 2 genes from the megakaryocyte lineage signature gene set were extracted from the myelofibrosis and control aggregate. 6684 myelofibrosis cells and 244 control cells were identified. *TGFB1*, the primary driver of bone marrow fibrosis, *LTBP1*, which binds the latent form of TGFβ and targets it for activation, and the proliferation marker *MKI67*, are all expressed in a higher % of MkP in myelofibrosis and also at higher levels per cell (blue dot indicates mean expression) than healthy donor controls. Percent of cells in which relevant gene is detected are shown below the x-axis. **5C: Left:** Louvain community detection based on the KNN weighted graph superimposed on a UMAP of the cells from the MF-MkP cluster demonstrates heterogeneity within MF-MkP. **Right**: MkP sub-clusters 4, 5 and 6 show high expression of the proliferation marker *MKI67*, not expressed by control MkP (supplementary), with high expression of the proliferation marker *TOP2A* also in cluster 4. *AURKA*, encoding the target of alisertib, a molecule currently in clinical studies for myelofibrosis (Gangat et al., 2017; Wen et al., 2015), is expressed by the minor sub-cluster 6. Pro-fibrotic cytokines *TGFB1* and *CXCL2* are expressed by the sub-clusters 1-6. 3 outlier cells are excluded from plots due to y-axis scaling for *CXCL2*. Blue dots on violin plot indicate mean level of expression. See also Suppl. Fig. S5.

Normal megakaryocytes have a low proliferation index and healthy donor MkP showed low expression of the proliferation marker *MKI67* (detected in only 6% of control MkP). By contrast, *MKI67* was detected in >30% of myelofibrosis MkP and the MkP cluster showed highest expression of *MKI67* among all myelofibrosis lineage clusters (Fig. 5B, Suppl. Fig. S5A) as well as enrichment of a G2M checkpoint gene signature (Suppl. Fig. S5B, Suppl. Table 7), suggesting that increased proliferation of MkP may contribute to the pathological accumulation of megakaryocytes in myelofibrosis, in addition to Mk-biased hematopoiesis.

### Myelofibrosis MkP demonstrate molecular heterogeneity with differential expression of proliferation and fibrosis genes

To identify distinct subpopulations of myelofibrotic MkP (MF-MkP), unsupervised clustering using Louvain community detection based on the KNN-weighted graph was performed on cells within the dominant Mk cluster (MkP2) from the myelofibrosis aggregate UMAP (Fig.2B, Suppl. Table 5). Seven sub-clusters were identified with distinct expression of fibrosis and proliferation-associated genes (Fig. 5C, Suppl. Fig. S5C). Genes encoding key mediators of fibrosis (*TGFB1* and *CXCL2)* were most highly expressed in MF-MkP clusters 1 – 5, whereas MF-MkP clusters 4 – 6 showed highest expression of proliferation markers *MKI67* and *TOP2A* and an G2M gene signature (Fig. 5C, Suppl. Fig. S5D). *AURKA* emerged as selectively expressed in clusters 6 and 4, with particularly high expression in the minor cluster 6 (Fig 5C). This is of interest as *AURKA* is the target for alisertib (MLN8237), recently demonstrated to promote megakaryocyte polyploidization and ameliorate the myelofibrosis phenotype in mouse models (Wen et al., 2015), with some efficacy also in patients with myelofibrosis (Gangat et al., 2019).

### Identifying myelofibrosis clone-specific cell surface targets

Increased expression of megakaryocyte genes in the myelofibrosis aggregate was noted to occur not just within the MkP cluster but also within clusters of uncommitted HSPCs and other lineage-affiliated clusters (Fig. 6A). This included intracellular proteins (*VWF* and *PF4*) and also cell surface antigens (*ITGAB1* [CD41] and *C6orf25 [G6B]*). Increased expression of *C6orf25,* encoding the G6B protein, was particularly striking (Fig. 6A). G6B is an immunoreceptor tyrosine-based inhibition motif (ITIM)-containing inhibitory immune receptor, considered to be exclusively expressed on mature megakaryocytes in normal hematopoiesis (Coxon et al., 2017; Senis et al., 2007). As the vast majority of healthy donor CD34^+^ lin^−^ HSPCs did not express megakaryocyte genes, and because mature megakaryocytes normally lose expression of CD34 during differentiation (Tomer, 2004), we hypothesized that aberrant co-expression of stem/progenitor and megakaryocyte surface antigens may enable selective identification of myelofibrosis clone-derived HSPCs.

**Figure 6:**
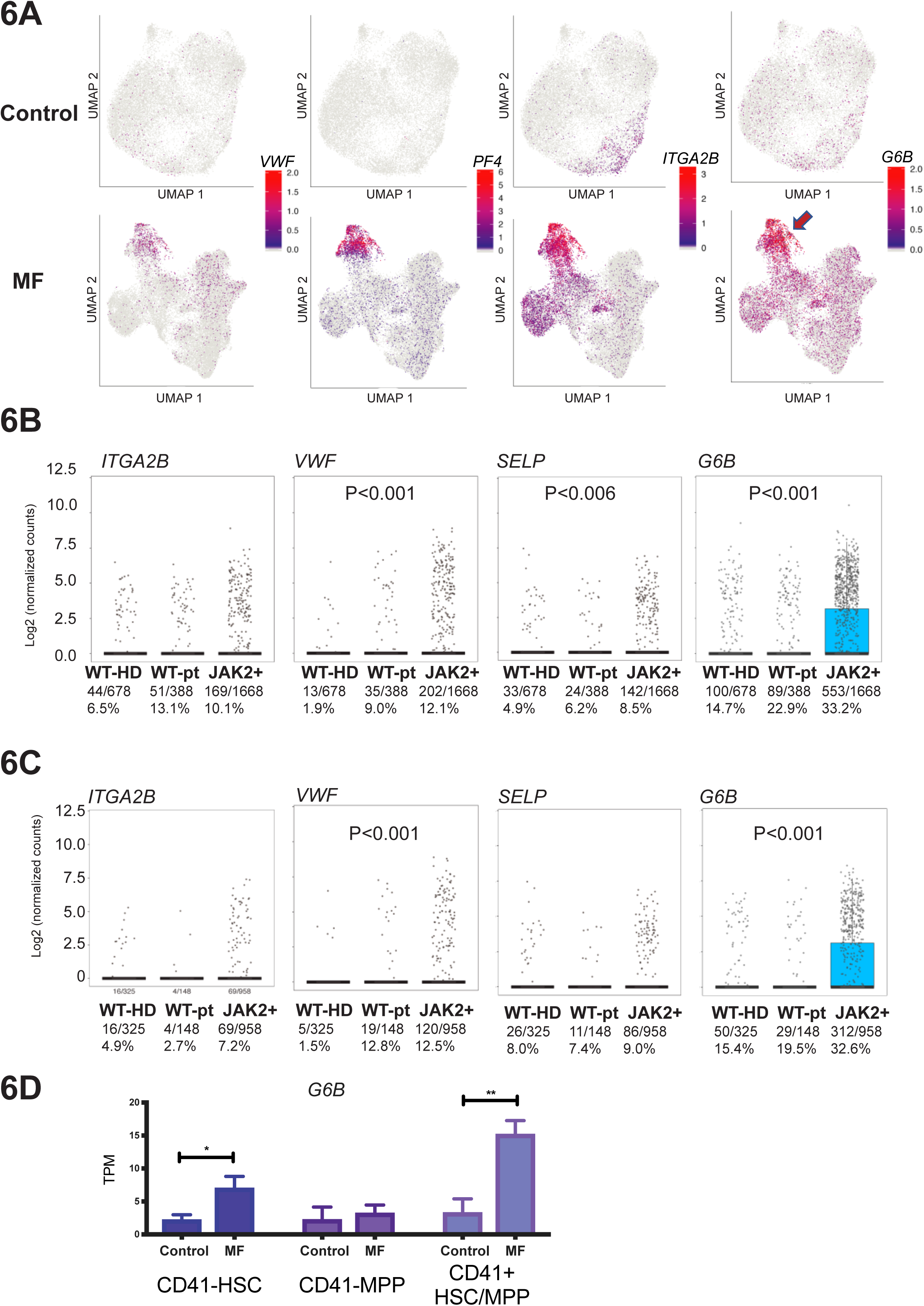
Increased expression of Mk-associated genes in myelofibrosis is not restricted to the MF-MkP cluster but is *JAK2V617F*+ mutant clone specific. **6A:** Increased expression of intracellular (*VWF, PF4*) and cell surface (*ITGA2B* [CD41], *G6B*) megakaryocyte genes is not limited to MF-MkP (arrow) in myelofibrosis, particularly for *G6B*. **6B:** Simultaneous targeted mutational profiling and RNA-sequencing (TARGET-Seq) of 2,734 individual CD34^+^ Lin^−^ HSPCs shows selective expression of megakaryocyte lineage genes *ITGA2B* (CD41), *VWF, SELP* and *G6B* in *JAK2V617F*-mutated and not wild-type cells within the same patients, or age-matched healthy donor control HSPCs. Fraction and % of cells in which gene expression was detected is shown. P-value for 3 × 3 chi-square is shown. **6C:** Expression of *ITGA2B* (CD41), *VWF, SELP* and *G6B* specifically in individual CD38-negative immunophenotypic stem cells (selected according to index sorting data) from healthy donors (normal), wild-type (WT) and *JAK2V617F*+ (JAK2+) myelofibrosis cells. **6D**: Expression of *G6B* in bulk-sorted control and myelofibrosis immunophenotypic HSC (CD34^+^lin^−^ CD38^−^CD45RA^−^CD90^+^), MPP (CD34+lin-CD38-CD45RA-CD90-) and CD41+ HSC/MPP (CD34^+^ lin^−^ CD38-CD45RA^−^ CD41^+^).TPM – transcripts per million. Chart shows mean ± SEM, n=4 for controls and n=3 for myelofibrosis; *P<0.05; **P≤0.01.

Patients with myelofibrosis have distinct genetic subclones of HSPCs, including residual wild-type (non-mutated) as well as clones with co-mutations in addition to driver mutations (*JAK2V617F* or *mutCALR*). To determine whether the increase in expression of megakaryocyte-associated genes was specific to mutant clone HSPCs or due to cell-extrinsic signals affecting both mutated and unmutated HSPCs, CD34^+^ lin^−^ HSPCs were analyzed by high-sensitivity mutational analysis and parallel transcriptome profiling (TARGET-Seq (Rodriguez-Meira et al., 2019)). 2734 cells were examined – 678 healthy donor cells plus 2056 myelofibrosis cells (388 *JAK2*-wild type and 1668 *JAK2V617F*-mutated). Expression of megakaryocyte genes, in particular *G6B*, was significantly higher in *JAK2V617F*-mutated HSPCs than in either wild-type cells from the same patients or in wild-type cells from healthy donors (32% vs 22.9% vs 14.7%, p<0.001, Fig. 6B). Wild-type cells from myelofibrosis patients also showed increased frequency of *G6B* expression, albeit to a much lower degree than *JAK2V617F*-positive cells, in keeping with cell-extrinsic signals also contributing to this aberrant megakaryocyte differentiation (Fig. 6B). The high-throughput TARGET-Seq and 10x Chromium datasets included all CD34^+^ lin^−^ cells. To determine if aberrant G6B expression was also present on the JAK2V617F+ stem cells as well as downstream progenitors, expression levels were verified in individual CD38-early stem/progenitor cells (HSC/MPP; Fig. 6C) and in 100-cell ‘mini-bulk’ preparations of FACS-isolated immunophenotypic CD34^+^ lin^−^ CD38^−^ CD45RA^−^ CD90^+^ HSCs, CD34^+^ lin^−^ CD38^−^ CD45RA^−^ CD90^−^ MPPs and CD41^+^ HSC/MPPs (Fig. 6D). Further, in two patients with 3+ co-mutations in addition to the driver *JAK2V617F* mutation, the increase in *G6B* was observed in all genetic sub-clones detected (Suppl. Fig. S6).

### Expression of cell surface G6B protein selectively identifies mutant clone derived HSPCs in myelofibrosis

High-throughput, single-cell proteomics by mass cytometry time of flight (CyTOF) was performed to enable simultaneous measurement of 20 surface proteins in multiple samples in parallel using barcode multiplexing (Fig. 7A, Suppl. Table 8). G6B was consistently detected at substantially higher levels in patients with primary and secondary myelofibrosis and with *JAK2V617F* and *mutCALR* driver mutations than in healthy donors (Fig. 7A, 7B). In addition, high cell surface G6B expression was also detected exclusively on *JAK2V617F*-mutated MPN cell lines (HEL, SET2) and not on the other leukemia cell lines K562, HL60, JURKAT and MARIMO and HEK human embryonic kidney cells (Suppl. Fig. S7). G6B expression was noted in both the CD41 positive and negative cell fractions in myelofibrosis by FACS (Fig. 7B).

**Figure 7:**
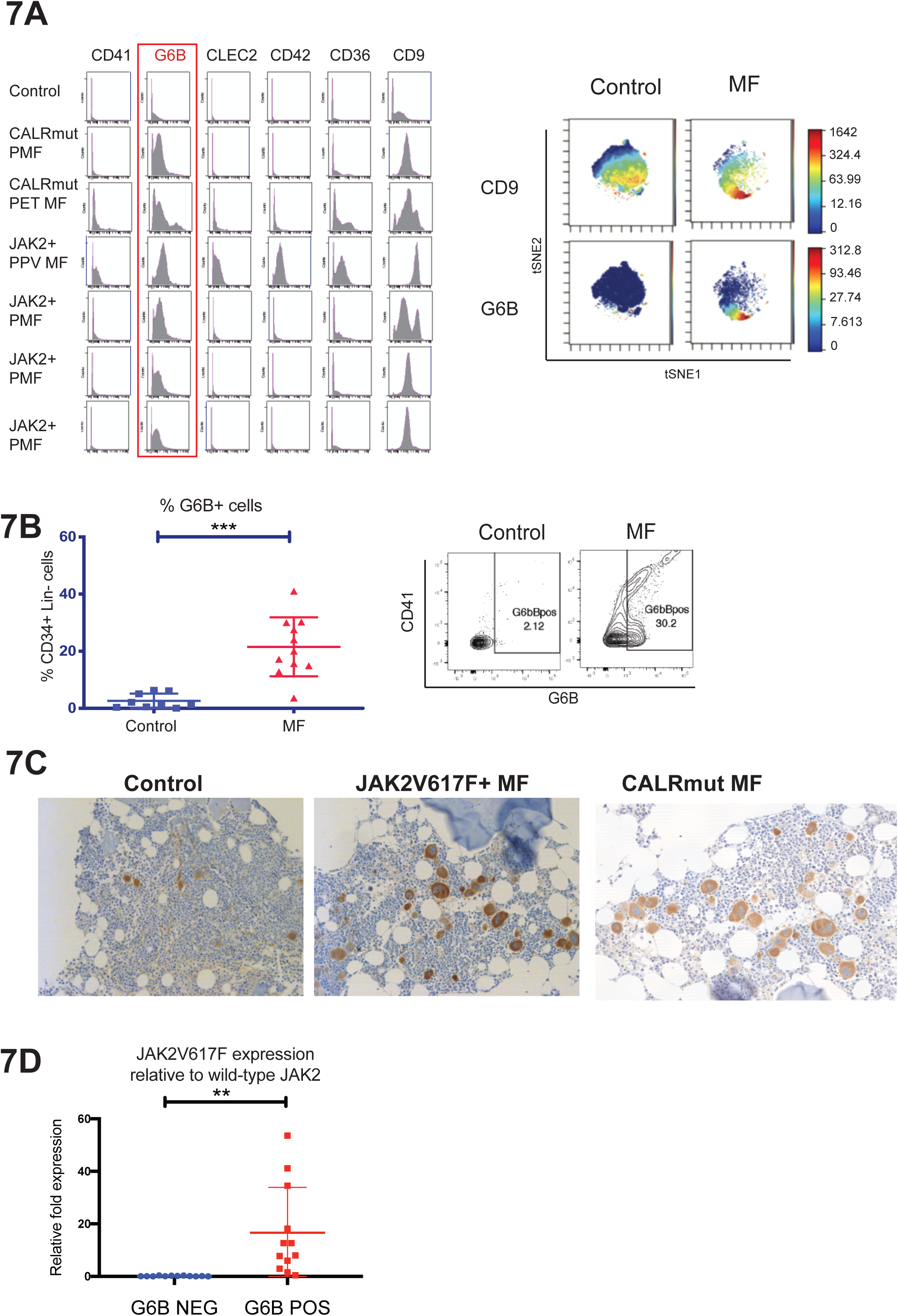
Expression of cell surface G6B, an immunoglobulin receptor, selectively identifies mutant clone-derived HSPCs in myelofibrosis. **7A: Left** - Expression of 6 megakaryocyte markers from a panel of 20 HSPC and megakaryocyte cell surface antigens assayed by time-of-flight mass spectrometry (CyTOF) shows distinct surface expression of G6B among HSPCs from patients with primary myelofibrosis (PMF) as well as myelofibrosis secondary to polycythaemia (PPV MF) and essential thrombocythemia (PET MF) with both JAK2V617F (JAK2+) and calreticulin (*mutCALR*) driver mutations. Histograms show cell count (y-axis) by expression level (x-axis). **Right** – viSNE dimensionality reduction plots on a representative control and myelofibrosis sample for CD9 and G6B, illustrating higher differential expression of G6B in myelofibrosis than control as compared to CD9. **7B: Left** - FACS analysis of G6B expression among CD34+ Lin-HSPCs showing significant increase in G6B+ cells in myelofibrosis (% GFP+ cells, 28.8 ± 5.5 *vs*. 2.4 ± 1.0); Chart shows mean+SEM, **P≤0.01 (t-test), controls (n=9); myelofibrosis (n=11). **Right –** Example FACS plot showing G6B cells are detected both in CD41+ and CD41-fractions. **7C:** Immunohistochemical staining for G6B (diaminobenzidine, DAB brown) of bone marrow trephine sections from healthy donor control and patients with JAK2V617F+ and mutCALR myelofibrosis showing marked expansion of G6B+ megakaryocytes and progenitors in myelofibrosis (6 cases studied). **7D:** Mononuclear cells from a healthy donor and a patient with *JAK2V617F*^+^ myelofibrosis were combined and 50 cell ‘minibulks’ sorted from the CD34^+^G6B^+^ and CD34+G6B-populations for Taqman qRT-PCR to quantify expression of *JAK2V617F* and wild-type *JAK2*. Chart shows *JAK2V617F* relative to *JAK2WT* expression for 12 minibulks from one representative experiment of 3 replicate experiments.

To examine G6B expression in bone marrow megakaryocytes *in situ,* immunohistochemical staining was performed on trephine biopsy sections from healthy donors and patients with *mutCALR* and *JAK2V617F*+ myelofibrosis, confirming expected expression on control megakaryocytes but with a dramatic increase in G6B+ cells in myelofibrosis (Fig. 7C).

Finally, to validate G6B as a potential target for therapies directed exclusively to mutant clone derived HSPCs, G6B positive and negative cells were FACS-isolated from healthy donor and myelofibrosis patient MNCs and expression of mutant vs. wild-type JAK2 determined by quantitative real time PCR (Moliterno et al., 2006). Strikingly, expression of mutant JAK2V617F was almost exclusively restricted to G6B positive cells (Fig. 7D). Together, these data identify G6B as a promising cell surface antigen to selectively target the aberrant megakaryocytic differentiation seen in myelofibrosis HSPCs.

## Discussion

Bone marrow transplant is currently the only potentially curative treatment for myelofibrosis, but is associated with significant risk and the vast majority of patients are ineligible due to age and comorbidities. The introduction of JAK inhibitors has led to significant improvement in symptomatic management, but the majority of patients continue to experience substantial morbidity and a significant reduction in life expectancy. New approaches to treatment are urgently required. Megakaryocytes are well recognized as the key cellular drivers of disease pathogenesis (Malara et al., 2018), however only one megakaryocyte-targeting therapy – alisertib, a specific inhibitor of aurora kinase – has been developed to date (Gangat et al., 2019; Wen et al., 2015). A major obstacle to identification of novel targets has been the inability to isolate bone marrow megakaryocytes from patients for detailed study. In the present study, we reasoned that aberrant megakaryopoiesis in myelofibrosis is very likely to be caused by aberrant differentiation of HSPCs, rather than proliferation of mature megakaryocytes, and that this process might be amenable to therapeutic targeting to ‘turn off the supply’. We therefore set out to characterise the distinct cellular and molecular features of megakaryocyte differentiation pathways in myelofibrosis, using a combination of single-cell approaches. We demonstrate an abnormal pathway for differentiation of megakaryocytes from uncommitted stem/progenitor cells in *JAK2V617F*-driven hematopoiesis, in addition to expansion of the normal shared trajectory between the erythroid and megakaryocyte lineages. Furthermore, a number of novel molecular targets that may inhibit the abnormal megakaryocyte differentiation and potentially ablate mutant clone-derived HSPCs and MkPs were identified.

Importantly, several key observations were only possible due to the single cell-level resolution of study, highlighting the power of single-cell technologies in understanding disease pathology and in novel therapeutic target discovery. Firstly, our data indicate that megakaryocyte-induced fibrosis is due to both a dramatic increase in MkP cell numbers, as well as increased production of fibrosis genes per cell, which are restricted to certain subpopulations of MkPs. Secondly, by simultaneously interrogating the mutational status and the transcriptome of individual cells, we demonstrated that certain megakaryocyte surface antigens, in particular G6B, are markedly over-expressed in mutant clone-derived HSPCs compared with wild-type HSPCs from myelofibrosis patients or healthy donor HSPCs. This validates combinatorial targeting of stem cell (e.g. CD34) and megakaryocyte (e.g. G6B) surface antigens e.g. with bispecific antibody therapies as a potential strategy worthy of further investigation for selective ablation of the myelofibrosis clone. As none of the currently available treatments for myeloproliferative neoplasms reliably induce clonal remissions or substantially reduce fibrosis, this work sets the stage for immunotherapeutic targeting of aberrant hematopoiesis in myelofibrosis. Furthermore, the approach we have adopted and the resulting insights are highly relevant to other studies seeking to identify cancer cell-specific drug targets and cancer-associated fibrosis in other malignancies, as well as non-malignant disorders of tissue fibrosis.

## Supporting information

Suppl. Item 1

## Acknowledgements

We thank all the patients who kindly donated samples, the MRC WIMM Flow Cytometry; Dr. Michalina Mazurczyk in the Mass Spectrometry Facility and Drs. Neil Ashley and Gemma Buck in the MRC WIMM Single Cell Facility; Dr. Alice Young in the NIH Intramural Sequencing Center; the NHGRI Flow Cytometry, and the National Institute for Health Research (NIHR) Oxford Biomedical Research Centre (BRC). This work was funded by a Wellcome Career Development Fellowship, an Academy of Medical Sciences Award and a L’Oreal-UNESCO Women in Science Award to B.P, a Medical Research Council (MRC) Senior Clinical Fellowship and CRUK Senior Cancer Research Fellowship to A.J.M., Bloodwise (project grant to B.P and A.J.M), a Cancer Research UK DPhil Prize Studentship to A.R-M., and the MRC Molecular Haematology Unit core award to A.J.M., and an MRC John Fell Fund award to B.P and A.J.M. The views expressed are those of the author(s) and not necessarily those of the NHS, the NIHR, the Department of Health, or the NIH.

## Author Contributions

B.P. designed, performed and analysed experiments, performed bioinformatic analyses and wrote the manuscript. G.W. designed, developed and performed bioinformatic analyses. A.R-M designed, performed and analysed experiments and performed bioinformatic analyses. E.H. performed and analysed experiments and contributed to bioinformatic analyses. R.L., J.O’S., N.S. performed experiments, processed samples and analysed data. S.A. performed experiments. Y.S. contributed the G6B antibody and interpreted data. O.W., D.R. and M.C. performed the immunohistochemical staining and analyses. D.I. performed experiments and assisted with protocol development. D.M. contributed clinical samples and data interpretation. I.R. and D.B. helped to supervise the project. S.T. designed, developed and supervised the computational data analysis. A.J.M. supervised the project, designed experiments, interpreted data and wrote the manuscript. All authors read and approved the submitted manuscript.

## Methods

### CONTACT FOR REAGENT AND RESOURCE SHARING

Further information and requests for resources or reagents will be fulfilled by

### Cell lines

HEL, JURKAT, K562, HEK, HL60 and MARIMO cells were obtained from the American Type Culture Collection (ATCC). SET2 cells were kindly provided by Dr. Jacqueline Boultwood and Dr. Andrea Pellagatti (Radcliffe Department of Medicine, University of Oxford). All cells were maintained in culture in RPMI-1630 supplemented with 10% fetal calf serum (FCS) and 1% penicillin-streptomycin. SET2 cells were supplemented with 20% FCS.

### Banking and processing of human samples

Patients and normal donors provided written informed consent in accordance with the Declaration of Helsinki for sample collection, tissue banking and use in research under either the INForMed Study, University of Oxford (IRAS: 199833; REC 16/LO/1376) or Imperial College London (approval reference: R13077; HTA licence 12275; REC 12/WA/0196). Cryopreserved peripheral blood mononuclear cells stored in FCS with 10% DMSO were thawed and processed by warming briefly at 37°C, gradual dilution into RPMI-1630 supplemented with 10% FCS and 0.1mg/mL DNAse I, centrifuged at 500G for 5 minutes and washed in FACS buffer (PBS + 2mM EDTA + 10% FCS).

### Fluorescent activated cell sorting (FACS) staining, analysis and cell isolation

FACS-sorting was performed using Becton Dickinson Aria III and cells isolated into 1.5ml Eppendorf tubes or 96-well plates depending on the experiment. Single color stained controls and fluorescence minus one (FMO) controls were used for all experiments. HSPCs were stained with the following antibody cocktail for 20 minutes at 4°C and passed through a 70 μm mesh cell strainer if necessary: CD34-APC-efluor780; Linege-BV510; CD38-PE-TxRed; CD123-PeCy7; CD45RA-PE; CD71-AF700; CD41-APC; CD90-BV421. The following antibody cocktail was used to analyse cell differentiation: CD34-APC-efluor780, CD71-AF700, CD36-FITC, CD41 PeCy7, CD42 PE, CD11b-APC, CD14-APC. 7AAD was used for live/dead cell exclusion. For G6B immunostaining, cells were stained with anti-human G6B (17-4) kindly supplied by Prof. Yotis Senis for 30 minutes at 4°C (1:100), washed and stained with goat anti-mouse Alexa Fluor 488 secondary IgG antibody (2:200 ThermoFisher Cat#A10680) for 20 minutes in the fridge and washed prior to staining with fluorescence-conjugated commercial antibodies.

### *In vitro* liquid culture differentiation assays

Cells were isolated by FACS into 1.5 μL eppendorfs, centrifuged at 500G for 5 minutes, resuspended in 100ul culture medium and plated in flat-bottom 96-well plates (Corning). Media used was Stemspan SFEM (StemCell Technologies #09650) + 1% Pen/Strep supplemented with recombinant human cytokines (Peprotech). Cells were analysed by FACS on days 6 and 14 (50 μl removed and replaced with fresh media).

### Cytospins and MGG

Cells were FACS-isolated into 1.5ml Eppendorf tubes, centrifuged and resuspended into 200 μl PBS and cytospun at 500RPM for 5 minutes onto Superfrost glass slides. May Grunewald Giemsa stain was prepared as per manufacturers protocol, filtered and slides stained in May-Grunewald for 7 minutes followed by 20 minutes in Giemsa then washed in distilled water, air dried and coverslip applied.

### Methocult assay

Single cells were FACS-isolated into flat bottomed 96-well plates containing 100 μl of MethoCult^TM^ H4435 Enriched (StemCell Technologies Cat#04435). Colonies were visually inspected and classified 11-14 days after plating. Lineage assignment was made by morphological assessment with verification of ambiguous colonies by plucking and FACS analysis.

### High-throughput single-cell RNA-sequencing (10X Chromium)

Cells were thawed, stained with FACS antibodies and sorted on an Aria III as described above and as per recommendations in the 10x Genomics Single Cell Protocols – Cell Preparation Guide. 15,000 CD34+ lineage negative cells were sorted into 20 μL PBS/0.05% BSA (non-acetylated) and then the cell number/volume adjusted to a target of 10,000 cells in 38 μL for loading onto the 10X Chromium Controller. Samples were processed according to the 10x protocol using the Chromium Single Cell 3’ library & Gel Bead Kit v2 (10x Genomics). In summary, cells and reagents were prepared and loaded onto the chip and into the Chromium Controller for droplet generation. RT was conducted in the droplets and cDNA recovered through demulsification and bead purification. Pre-amplified cDNA was used for library preparation, multiplexed and run on a MiSeq using MiSeq Nano Reagent Kit V2 (Illumina Cat#102-2001). CellRanger was used to estimate the number of cells, and samples then sequenced on a HiSeq 2500 using v4 chemistry to obtain 40-50,000 reads per cell.

### TARGET-Seq

High-sensitivity single cell mutation analysis and parallel RNA-sequencing was performed as previously described (Rodriguez-Meira et al., 2019). Counts were downloaded from <u>GSE122198</u>, normalized by library size and log2-transformed as previously described (Rodriguez-Meira et al., 2019). Cells were classified into WT-normal (cells from normal donors), WT-patient (non-mutant cells present in patient samples) and mutant (cells from patient samples carrying mutations in the genes targeted).

### RNA sequencing of ‘mini-bulk’ HSPC populations

100 cells from each population were isolated by FACS into 4 μl of lysis buffer containing oligo-dT primer and dNTP mix in 0.2 mL PCR tubes. Cell lysis, RT and PCR preamplification and purification was performed using the Smart-Seq 2 protocol as previously published (Picelli et al., 2014). Libraries were pooled and tagmentation performed using the Illumina Nextera XT DNA sample preparation kit (Illumina Cat #FC-131-1024), libraries pooled and sequenced on a HiSeq 2000.

### Antibody labelling with metal conjugates and mass cytometry (CyTOF)

Antibodies were purchased pre-conjugated when commercially available. Non-available antibodies were conjugated to lanthanide metals using Maxpar X8 antibody labelling kit according to the manufacturer protocol (version 10). The antibody cocktail used is listed in Suppl. Table S8. For barcoding and staining, cells were washed with Maxpar PBS buffer (Fluidigm #201058) and stained with 0.5 μM Cell-ID Cisplatin Viability Stain (Fluidigm #201064) in 200 μL Maxpar PBS for 5 mintutes at room temperature for dead cell exclusion. The reaction was quenched with Maxpar Cell Staining Buffer (CSB, Fluidigm #201063) and cells fixed, permeabilized and barcoded using the Cell-ID 20-Plex Pd Barcoding Kit (Fluidigm #201060) as per the manufacturers user guide. Barcoded cells were washed, combined and stained with the antibody cocktail as per Suppl. Table 8 for 30 minutes at room temperature. Cells were washed with Maxpar Cell Staining Buffer (Fluidigm #201068), fixed in 1.6% formaldehyde, washed and resuspended in Fix&Perm Buffer (Fluidigm Cat#201067) with Cell-ID intercalator-Ir (Fluidigm #201103B) and incubated overnight at 4°C. The following day, cells were washed and analysed on a Helios (Fluidigm). The mass cytometer was tuned and QC was run prior to acquiring samples according to the manufacturers’ recommendations.

### G6B Immunohistochemistry

Sections of formalin fixed and paraffin embedded (FFPE) bone marrow trephine biopsies were processed as follows: paraffin removed, antigen retrieval performed using citrate (Roche Cell Conditioning 2 Cat#950-123) pre-treatment for 30 minutes, washed and incubated with G6B antibody diluted 1:100 in Ventana’s DISCOVERY antibody diluent (Roche Cat#760-108) for 60 minutes at room temperature. Secondary detection was performed using UltraMap DAB anti-Ms HRP detection kit (Roche #760-152) for 16 minutes and slides counterstained with hematoxylin (Roche #760-2021) for 4 minutes and Bluing reagent (Roche #760-2037) for 4 minutes.

### Sorting G6B+ and G6B-HSPCs for JAK2V617F qRT-PCR

For each experiment, MNCs from myelofibrosis patients and healthy donor controls were thawed and combined 1:1 in FACS buffer prior to antibody staining as described above. 50 G6B+ and G6B-cells were sorted into each well of a 96-well PCR plate (10 replicates per population for each experiment), containing CellsDirect One-Step qRT-PCR kit 2X Reaction Buffer and SuperScript III RT/Platinum Taq Mix (Thermo Fisher Cat#11753100), Ambion SUPERase-In RNase inhibitor (Thermo Fisher Cat#AM2694), TE buffer, JAK2 forward and reverse primers and wild-type and JAK2V617F-specific probe mix (see Key Resources Table). RT and PCR were performed as per the kit protocol with 18 pre-amplification cycles then diluted 5x in TE buffer. Taqman RT-PCR was performed in a 20 μL reaction volume using 4 μL of the diluted cDNA, Taqman Fast Advance Mastermix (Thermo Fisher Cat#4444556) and the primers/probes as detailed in the Key Resources Table. Custom Taqman assays were designed as previously described (Moliterno et al., 2006) using RT-PCR primers flanking the mutant region plus two Taqman PCR probes specific for the normal or mutant sequence. An Applied Biosystems 7500 Fast Real-Type PCR system was used with the default PCR conditions, with each replicate run in duplicate. Intra-assay replicates varying more than 5% were excluded.

### 10x Genomics single-cell RNA sequencing data pre-processing

Sequencing data in the binary base call (BCL) format were demultiplexed. Unique molecular identifier (UMI) counts for given genes were obtained by aligning FASTQ files to the human reference genome (GRCh38) using Cell Ranger software (version 2.0.0) from 10x Genomics. CellRanger “count” pipeline results from each of individual libraries from three patients and two healthy donors were then aggregated using default parameters to generate a gene count matrix based on CellRanger “aggr” standard pipeline. The UMI counts (> 1,000 and ≤ limited maximum UMIs), the number of detected genes (> 500 and ≤ limited maximum number of detected genes) and the percentage of mitochondrial gene expression (< 10%) per cell used as the cut-off criteria described in Suppl. Table 3. Following these filters, 47,804 cells passed quality control for the whole sample aggregation. We excluded 617 out of 48,421 cells from further analyses. We scaled UMI counts by the total library size multiplied by 10,000. The normalized expression values were then log transformed. We regressed out the unwanted source of variation (library size, percentage of mitochondrial genes and batches from patients and healthy donors) from gene expression values by applying the linear regression model using the limma package (Ritchie et al., 2015).

### Bioinformatics Analysis and R Code

Methods under submission; can be requested from

### Quantification and Statistical Analysis

#### Flow cytometry and CyTOF data analysis

Flow cytometry data was analysed using FlowJo software (v10.5.3). Summary charts and associated statistical analyses were performed using GraphPad Prism (v8.1.0). Helios CyTOF Software (v6.7) was used for processing of FCS 3.0 files, normalization to EQ Beads, concatenation of multiple files and debarcoding. Data was then analysed and histograms and viSNE plots generated using CytoBank.org.

Statistical tests used, numbers of replicates and definitions of statistical significance are described in the relevant figure legends. All bar charts show mean ± standard error of the mean and were generated using GraphPad Prism (v.8.1.0).

### Data and software availability

Data has been submitted to GEO (Accession Number will be provided on publication). TARGET-Seq single cell RNA-sequencing data is available at GEO: GSE105454. The Shiny application for visualisation of the data from patients and healthy donors in this study is available at https://github.com/supatt-lab/SingCellR-myelofibrosis.

### Supplemental Information

**Suppl. Figure 1, Relating to Figures 1 and 2. Isolation and characterization of CD41+ and CD41-CD34+lin-CD38-CD45RA-early stem/progenitor cells, distribution of individual donor cells in sample aggregates and proportions of cells classified within each lineage-affiliated cluster.**

**S1A.** Sort strategy for isolation of Lin-CD34+CD38-CD45RA-CD41+ (CD41+ HSC/MPP) and Lin-CD34+CD38+CD45RA-CD41+ (CD41+ CMP/MEP) cells. Example plot for healthy donor control (control) and myelofibrosis (MF) shown.

**S1B.** Lin-CD34+CD38-CD45RA-CD41+ (CD41+ HSC/MPP) and Lin-CD34+CD38+CD45RA-CD41+ (CD41+

CMP/MEP) compartments contain cells with typical blast cell morphology and not mature megakaryocytes. Representative cells shown, cells isolated by FACS and stained with May Grunewald Giemsa. 100X magnification.

**S1C.** Single-cell colony output from control and myelofibrosis CD41-HSC, CD41-MPP and CD41+ HSC/MPP cells FACS-isolated into individual wells of 96-well plates containing methylcellulose. Total colonies counted = 250 sorted from healthy control (n=5) and myelofibrosis (n=6) donors.

**S1D:** Distribution of cells from individual healthy donor (left) and myelofibrosis (right) donors over UMAP plots of patient/control aggregates after batch correction.

**S1E:** % of total cells contained within undifferentiated hematopoietic stem/progenitor cell (HSPC), erythroid (Ery), lymphoid (Lym), myeloid (Mye) and megakaryocyte (Mk) clusters for the control and myelofibrosis aggregates (as classified in Figure 2A).

**Suppl. Fig. 2, Relating to Figure 2. Single-gene UMAP plots for healthy donor and control aggregates illustrating classification of clusters**

Expression of three individual genes from each of the lineage signature gene sets for cells from healthy donor controls (**2A**) and myelofibrosis patients (**2B**). Cells on Uniform Manifold Approximation and Projection (UMAP) plots are colored according to expression levels for each gene from not detected (grey) → low (blue) → high (red). Violin plots show expression levels for each gene for cells classified into 8 k-means clusters (see Figure 2A). Violin plots: x-axis – cluster ID. For controls, cluster 1 – HSPC 1; cluster 2 – HSPC 2; cluster 3 – HSPC 3; cluster 4 – myeloid; cluster 5 – HSPC 4; cluster 6 – erythroid; cluster 7 – erythroid/megakaryocyte; cluster 8 – lymphoid. For myelofibrosis, cluster 1 – HSPC 1; cluster 2 – HSPC 2; cluster 3 – HSPC 3; cluster 4 – erythroid; cluster 5 – lymphoid; cluster 6 – myeloid; cluster 7 – megakaryocyte progenitor 1; cluster 8 – megakaryocyte progenitor 2 (see also Suppl. Tables 4 and 5). Y-axis - log(Normalized UMI). Abbreviations - Mye - myeloid; Lym - lymphoid; Ery – erythoid; MK – megakaryocyte.

**Suppl. Fig. 3, Relating to Figure 4. Potential molecular regulators of erythroid differentiation.**

Heatmap showing expression of 51 transcription factors selected from those showing progressively increased/decreased expression along the trajectory path from the earliest undifferentiated HSPC cluster (green) to the erythroid cluster (blue) in the aggregate of all healthy donor and myelofibrosis cells. Color legend shows log (Normalized UMI) expression level from low (blue) to high (red).

**Suppl. Fig. 4, Relating to Figure 4. Potential molecular regulators of aberrant megakaryocyte differentiation.**

Heatmap showing expression of 51 transcription factors selected from those showing progressively increased/decreased expression along the trajectory path from the earliest undifferentiated HSPC cluster (green) to the megakaryocyte cluster (purple). The path was selected from the aggregate of all healthy donor and myelofibrosis cells, but the megakaryocyte cluster almost exclusively (>99%) comprises myelofibrosis cells. Color legend shows log(Normalized UMI) expression level from low (blue) to high (red).

**Suppl. Fig. 5, Relating to Figure 5. Myelofibrosis megakaryocyte progenitors (MF-Mk) highly express marker genes of proliferation and are heterogeneous, with proliferative and pro-fibrotic subpopulations.**

**S5A:** Expression of the proliferation marker gene *MKI67* is highly expressed almost exclusively in the MF-MkP cluster.

**S5B:** Gene set enrichment analysis of genes differentially expressed in myelofibrosis megakaryocytes (MF-Mk2 cluster, see also Figure 2A) *vs.* all other myelofibrosis HSPC clusters shows significant enrichment of G2M checkpoint genes. See also Suppl. Table 7.

**S5C:** Heatmap of top 10 differentially expressed genes in the seven MF-MkP subclusters of the myelofibrosis Mk2 cluster (see also Fig.2A).

**S5D:** UMAP showing expression of G2M and S phase cell cycle genes over the MF-MkP subclusters.

**Suppl. Fig. 6, Relating to Figure 6. *G6B* expression within distinct molecular subclones.**

Expression of *G6B* is detected in all genetic subclones in two myelofibrosis patients with 3+ mutational subclones. x-axis – number of cells in which *G6B* expression was detected out of all cells of each molecular subclone studied.

**Suppl. Fig. 7, Relating to Figure 7. Expression of *G6B* is detected only in the *JAK2V617F*-mutated cell lines HEL and SET2 and not in *JAK2* wild-type leukemia cell lines**.

MARIMO (mutCALR acute myeloid leukemia), HL60 (acute myeloid leukemia), JURKAT (T-cell leukemia), K562 (chronic myeloid leukemia), or human embryonic kidney HEK cells.

### Supplementary Tables

**Suppl. Table 1, Relating to Figures 1-8:** Clinical details of all myelofibrosis patients studied and healthy donor controls. Abbreviations: MF – myelofibrosis; Con – healthy donor controls; Hydroxy – hydroxycarbamide; EPO – recombinant human erythropoietin. DIPPS – dynamic international prognostic scoring system (Passamonti et al., 2010).

**Suppl. Table 2 (See excel file), Relating to Figures 2 - 5**: Detailed clinical information of patients from whom samples were studied by high-throughput single-cell RNA-sequencing (10x Chromium).

**Suppl. Table 3, Relating to Figures 2 – 5:** Quality control and cells filtered out during quality control steps on single cell RNA-sequencing data. Abbreviations: HD_Agg – aggregate of all healthy donor cells; MF_Agg – aggregate of all myelofibrosis cells; All_Agg – aggregate of all healthy donor plus myelofibrosis cells.

**Suppl. Table 4 (see excel file), Relating to Figure 2**: Genes differentially expressed in the 8 k-means clusters for the healthy donor aggregate (up to 50 genes listed per cluster).

**Suppl. Table 5 (see excel file), Relating to Figure 2**: Genes differentially expressed in the 8 k-means clusters for the myelofibrosis donor aggregate (up to 50 genes listed per cluster).

**Suppl. Table 6, Relating to Figures 2, 3 and 5**. Genes included in HSPC lineage and fibrosis signature gene sets and G2M and S phase gene sets.

**Suppl. Table 7, Relating to Figure 5 (see excel file).** Gene set enrichment analysis of genes differentially expressed in myelofibrosis megakaryocytes (MkP 2 cluster).

**Suppl. Table 8, Relating to Figure 7**. Antibodies used for cell surface antigens in CyTOF panel.

### Other Suppl. Items

**Suppl. Item 1, Relating to Fig. 4.** .html file containing 3-dimensional k-nearest neighbor (KNN) graph aggregate of all 47,804 cells from healthy donors and patients with myelofibrosis with cells colored according to unsupervised k-means clustering.

## Supplementary Tables

**Suppl. Table 1.** Demographic and clinical details of patients and healthy donors studied.

| Group | Study ID | Age | Sex | Diagnosis | Known mutations | Treatment | WHO Fibrosis in BM | DIPPS risk group | Figure data contributing to |
| --- | --- | --- | --- | --- | --- | --- | --- | --- | --- |
| **MF** | 001/001 | 70 | M | PMF | JAK2V617F, CBL, SRSF2 | Nil | 3 | Int-1 | 1A, 7B, 6B, 6C, S7 |
|  | 001/002 | 56 | M | PET MF | CALR type 2, U2AF1 | Ruxolitinib | 3 | Int-2 | 1A, 7B, 6B, 6C |
|  | 001/020 | 61 | M | PET MF | CALR Type 1 | Ruxolitinib | 3 | Low | 1A |
|  | 001/023 | 86 | M | PMF | JAK2V617F, ASXL1, TET2, U2AF1 | Ruxolitinib | 3 | Int-2 | 1A, 6B, 6C, 7B |
|  | 001/027 | 40 | M | PET MF | JAK2V617F | Nil | 3 | Low | 1A |
|  | 001/037 | 59 | F | PMF | JAK2V617F | Nil | 3-4 | High | 6B, 6C, S7 |
|  | 001/038 | 60 | M | PPV MF | JAK2V617F | Hydroxy | 3 | Int-1 | 1A, 6B, 6C, 7A, |
|  | 001/040 | 73 | F | PPV MF | JAK2V617F | Ruxolitinib | 3 | Int-2 | 1A, 1E, 6B, 6C, 7A, 7B, 7D |
|  | 001/042 | 57 | F | PET MF | MPL TRP515LYS | Nil | 2-3 | Int-1 | 7B |
|  | 001/047 | 81 | F | PMF | CALR Type 1 | Ruxolitinib | 3 | Int-1 | 1A, 1E, 7A, 7B |
|  | 001/054 | 76 | F | PET MF | JAK2V617F, ETV6 | Nil (prior anagrelide) | 3 | Int-2 | 1A, 7D |
|  | 001/055 | 60 | F | PET MF | JAK2V617F | Anagrelide | 3 | Int-1 | 6B, 6C |
|  | 001/057 | 67 | F | PPV MF | JAK2V617F | Ruxolitinib | 2 | Int-1 | 1A, 6B, 6C, 7D |
|  | 001/076 | 70 | F | PET MF | CALR Type 1 | Luspatercept | 2 | Int-1 | 7B |
|  | 001/083 | 52 | F | PET MF | CALR Type 1, ASXL1 | Ruxolitinib, EPO, post-HCT | 3 | Int-2 | 7B |
|  | 001/098 | 58 | F | PMF | CALR Type 2, ASXL1 | Nil | 2-3 | Int-1 | 7B |
|  | 001/114 | 67 | M | PPV MF | JAK2V617F | Hydroxy | 2-3 | Int-1 | 7B |
|  | 006/002 | 67 | F | PMF | JAK2V617F | Momelotinib | 3 | Int-1 | 1A, 6B, 6C, 7B |
|  | 010/022 | 65 | F | PPV MF | JAK2V617F | Hydroxy | 2 | Int-1 | 1A, 1C, 1D, 2, 3, 4, 5, 6A, 6D |
|  | 010/054 | 77 | M | PPV MF | JAK2V617F | Ruxolitinib | 3 | Int-2 | 1A, 1C, 1D,<br>2, 3, 4, 5,<br>6A, 6D |
|  | 010/047 | 56 | F | PMF | JAK2V617F | Ruxolitinib | 3 | Int-2 | 1A |
|  | 010/005 | 62 | M | PMF | JAK2V617F | Danazol | 3 | Int-2 | 1A, 7A |
|  | 010/027 | 72 | M | PET MF | JAK2V617F | Ruxolitinib,<br>EPO | 2 | Int-2 | 1A |
|  | 010/028 | 74 | M | PPV MF | JAK2V617F | Ruxolitinib | 2 | Int-1 | 1A |
|  | 010/003 | 79 | M | PPV M | JAK2V617F | Ruxolitinib | 3 | Int-2 | 1A, 1C, 1D,<br>2, 3, 4, 5,<br>6A, 6D |
|  | 001/055 | 54 | F | PMF | JAK2V617F | Ruxolitinib | 2 | Int-1 | 6C |
|  | 001/019 | 78 | M | PET MF | JAK2V617F | Ruxolitinib,<br>EPO | 3 | Int-2 | 6C |
|  | 001/056 | 60 | M | PMF | JAK2V617F | Nil | 3 | Int-1 | 6C,S7 |
| **CON** | 009/001 | 63 |  |  |  |  |  |  | 1A, 7A, 7B |
|  | 009/002 | 42 |  |  |  |  |  |  | 1A,1E, 7A |
|  | 009/003 | 35 |  |  |  |  |  |  | 1A |
|  | 009/005 | 60 |  |  |  |  |  |  | 1A, 1E, 7A |
|  | 014/001 | 44 |  |  |  |  |  |  | 1A, 7B, S1C |
|  | 014/002 | 26 |  |  |  |  |  |  | 1C, 1D, S1C |
|  | 014/003 | 32 |  |  |  |  |  |  | 1A, S1C |
|  | 010/9001 | 54 |  |  |  |  |  |  | 1A, 1C, 1D,<br>7A, 7B S1C |
|  | 010/9002 | 58 |  |  |  |  |  |  | 1A, 7B, S1C |
|  | 011/001 | 40 |  |  |  |  |  |  | 1A, 7A |
|  | 012/9001 | 44 |  |  |  |  |  |  | 1A, 1C, 1D,<br>1E |
|  | 012/9002 | 53 |  |  |  |  |  |  | 1A, 2, 3, 4,<br>5, 6A, 6D |
|  | 012/9003 | 41 |  |  |  |  |  |  | 1A, 1E |
|  | 012/9004 | 33 |  |  |  |  |  |  | 1A, 1C, 1D |
|  | 012/9005 | 28 |  |  |  |  |  |  | 1A |
|  | 012/9006 | 51 |  |  |  |  |  |  | 1A, 2, 3, 4,<br>5, 6A, 6D |
|  | 012/9007 | 36 |  |  |  |  |  |  | 1A |
|  | 007/002 | 61 |  |  |  |  |  |  | 7D |
|  | 007/003 | 78 |  |  |  |  |  |  | 7D |
|  | 007/005 | 73 |  |  |  |  |  |  | 7D |

**Supplementary Table 3.** QC and Filtering of 10X scRNAseq data.

|  |  | **MF_aggregated** | **HD_aggregated** | **All_aggregated** |
| --- | --- | --- | --- | --- |
| **filtering cutoff** | maximum UMIs | 30000 | 23000 | 30000 |
|  | minimum UMIs | 1000 | 1000 | 1000 |
|  | % of mitochondrial genes | < 10% | < 10% | < 10% |
|  | maximum detected genes | 5000 | 4200 | 5000 |
|  | minimum detected genes | 500 | 500 | 500 |
| **Total cell number sequenced** |  | 30088 | 18333 | 48421 |
| **Cells included in analysis** |  | **29536** | **18249** | **47804** |
| **Average detected genes per cell** |  | **2119** | **1812** | **1998** |
| **The number of highly variable genes** |  | **1095** | **1050** | **841** |

**Supplementary Table 6.**
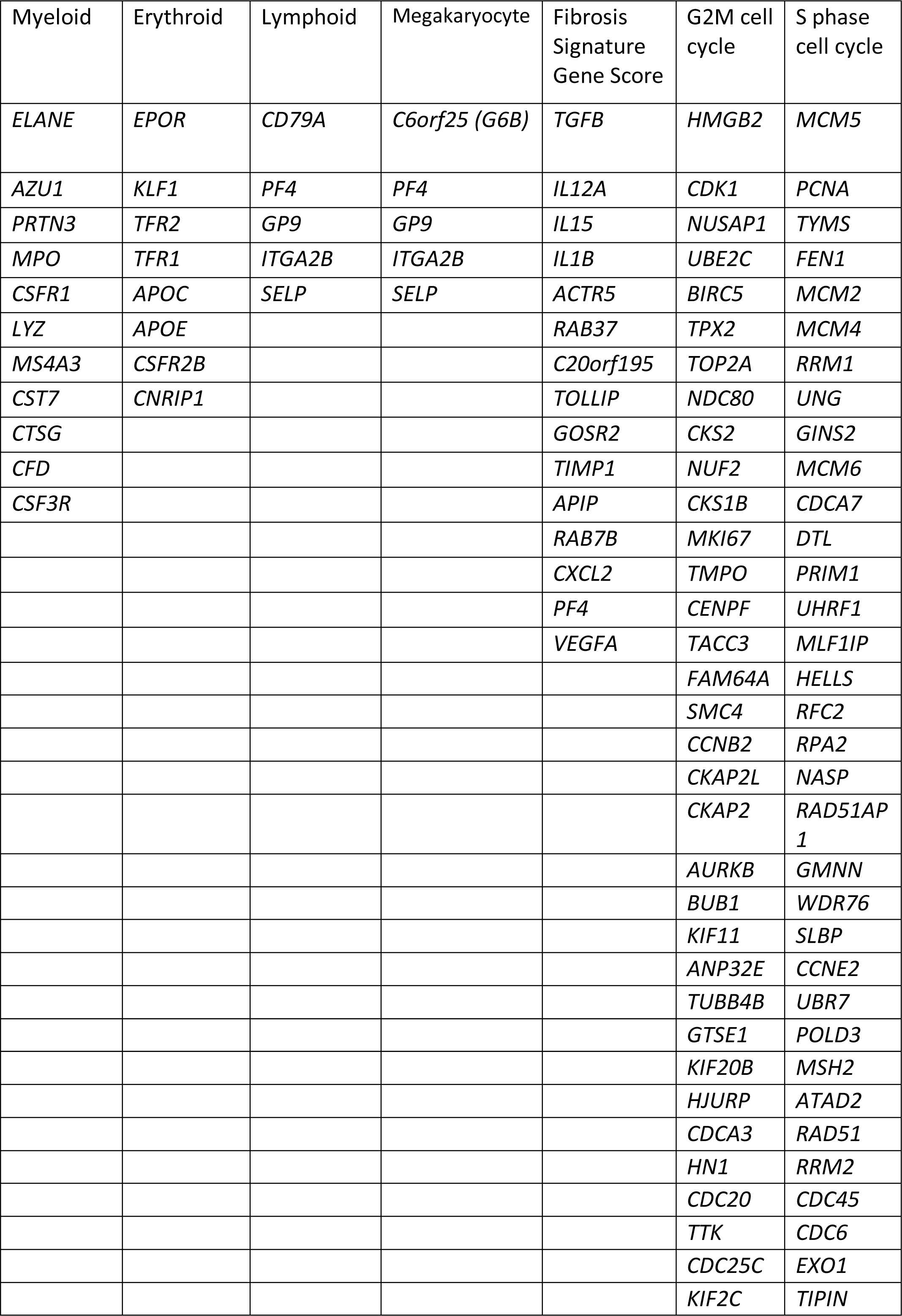
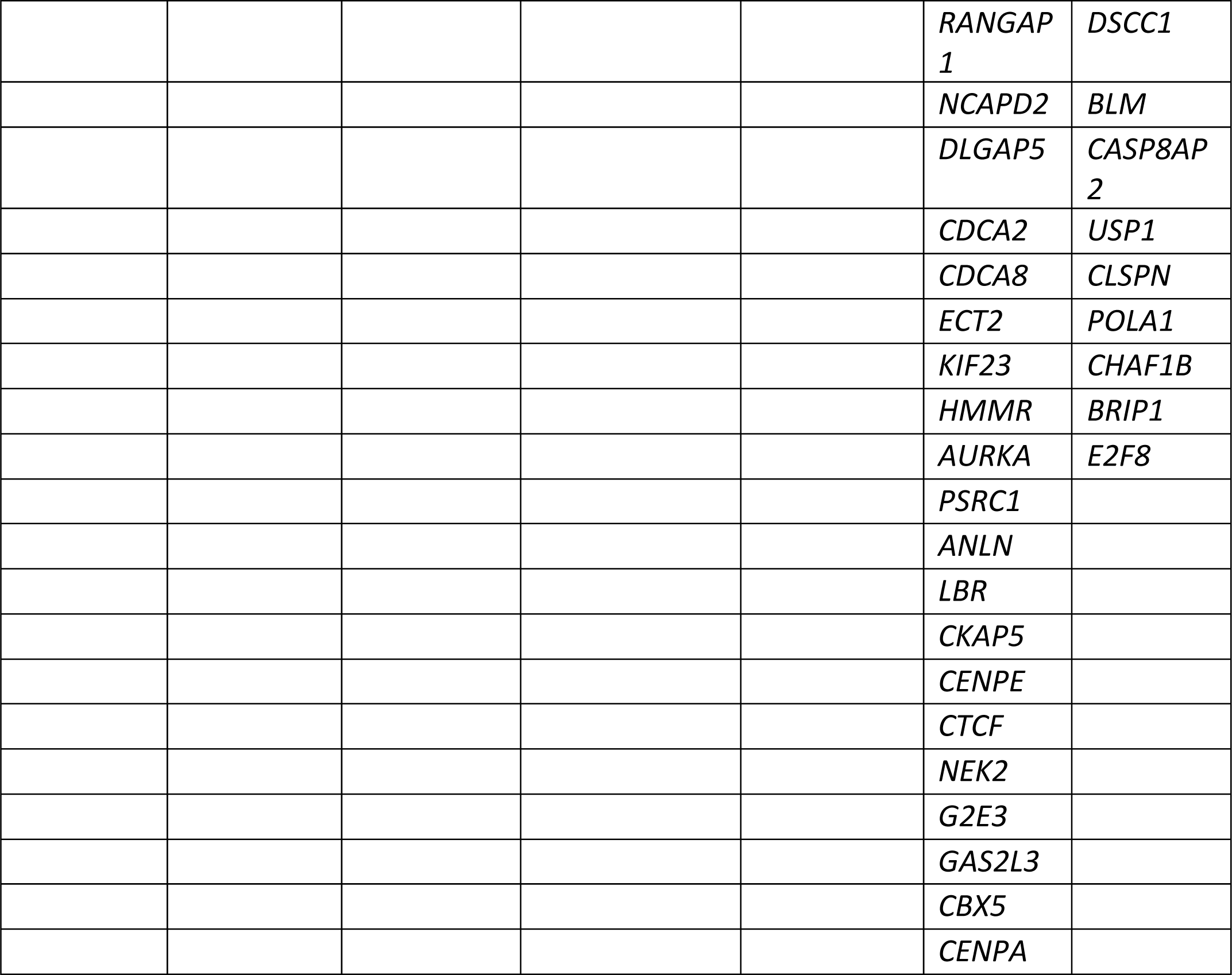
Genes included in signature gene sets.

**Supplementary Table 8.** Antibody panel used for CyTOF.

| **Marker** | **Clone** | **Supplier** | **Catalogue number** | **Titration (uL/100ul)** | **Isotope** |
| --- | --- | --- | --- | --- | --- |
| CD41 | HIP8 | Fluidigm | 3089004B | 1.0 | 89Y |
| CD19 | SJ25-C1 | Invitrogen | MHCD1906 | 0.2 | 111Cd |
| CD9 | HI9a | Biologend | 312102-B227707 | 1.0 | 142 Nd |
| CD45RA | HI100 | Fluidigm | 3143006B | 1.0 | 143 Nd |
| CD42b | HIP1 | Fluidigm | 3144020B | 1.0 | 144 Nd |
| CD4 | RPA-T4 | Fluidigm | 300502 | 0.5 | 145 Nd |
| CD34 | 581 | Fluidigm | 3148001B | 1.0 | 148 Nd |
| CD56 | NCAM16.3 | Fluidigm | 3149021B | 1.0 | 149 Sm |
| CD123 (IL-3R) | 6H6 | Fluidigm | 3151001B | 1.0 | 151 Eu |
| CD3 | UCHT1 | Fluidigm | 3154003B | 0.5 | 154 Sm |
| CD36 | 5-271 | Fluidigm | 3155012B | 1.0 | 155 Gd |
| CD14 | M5E2 | Biologend | 301802 | 0.3 | 160 Gd |
| CD90 | 5.00E+10 | Fluidigm | 3161009B | 1.0 | 161 Dy |
| CD49f | G0H3 | Fluidigm | 3164006B | 1.0 | 164 Dy |
| CD44 | BJ18 | Fluidigm | 3166001B | 1.0 | 166 Er |
| RXFP1 | 933344 | R&D Systems | MAB8898 | 1.0 | 170 Er |
| CD38 | HIT2 | Fluidigm | 3172007B | 1.0 | 172 Yb |
| CLEC2 | AYP1 | Biologend | 372002 | 1.0 | 174 Yb |
| CD71 | OKT-9 | Fluidigm | 3175011B | 1.0 | 175 Lu |
| G6B | clone 17-4 | Collaborator | NA | 2.0 | 176 Yb |

