## Supplementary figures and images for "Single-cell analyses reveal aberrant pathways for megakaryocyte-biased hematopoiesis in myelofibrosis and identify mutant clone-specific targets"

### Supplemental Figures

## S1A

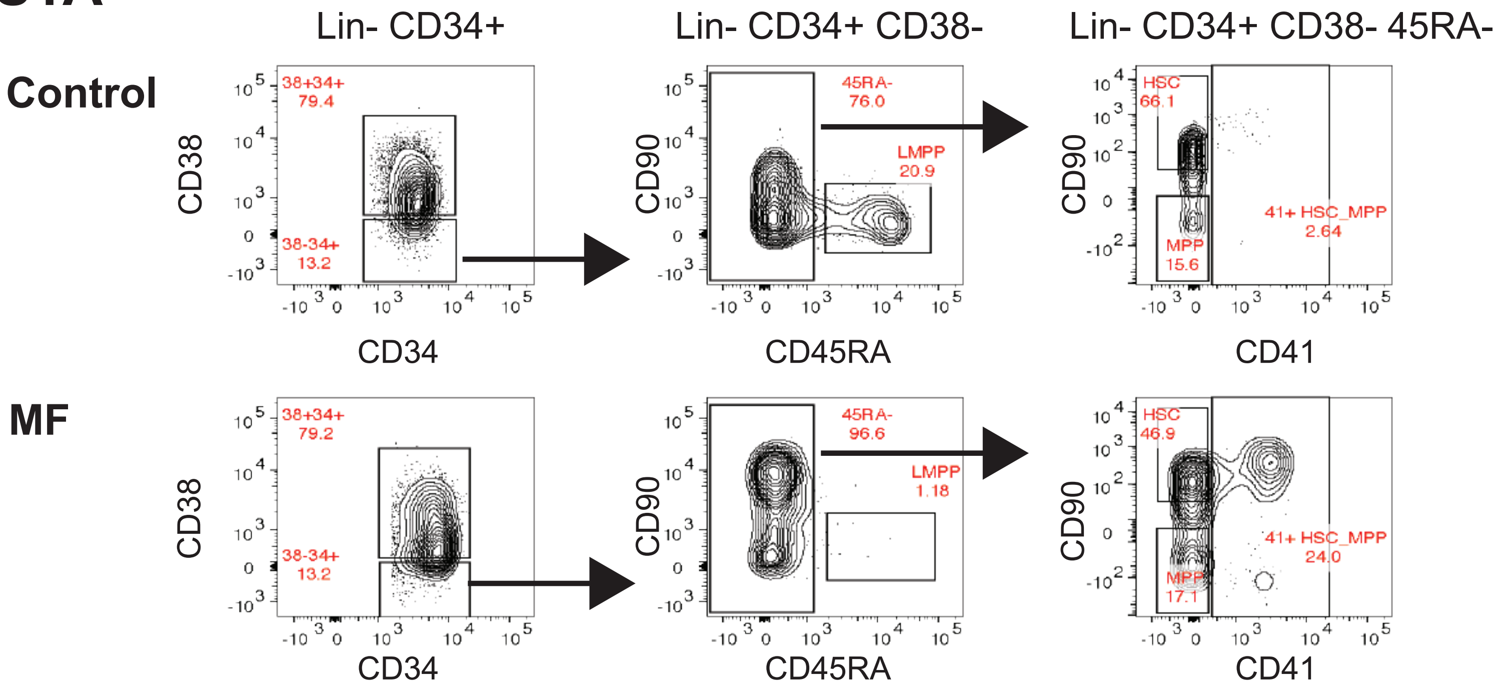


## S1B

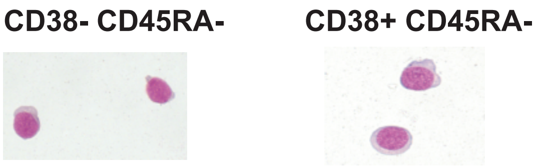


## S1C

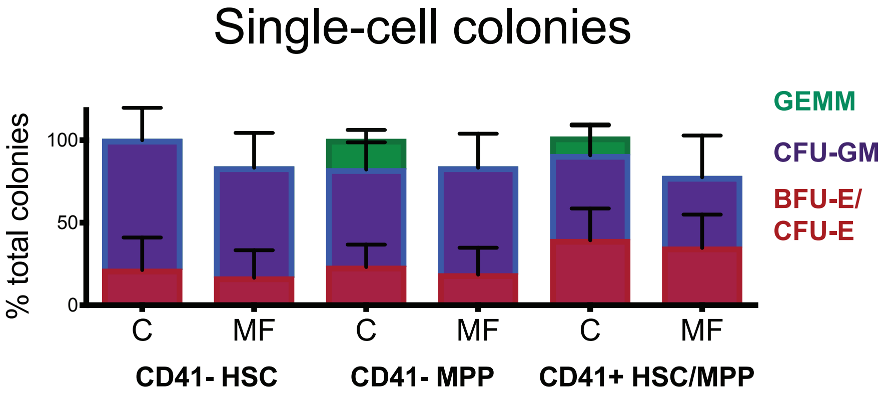


## S1D

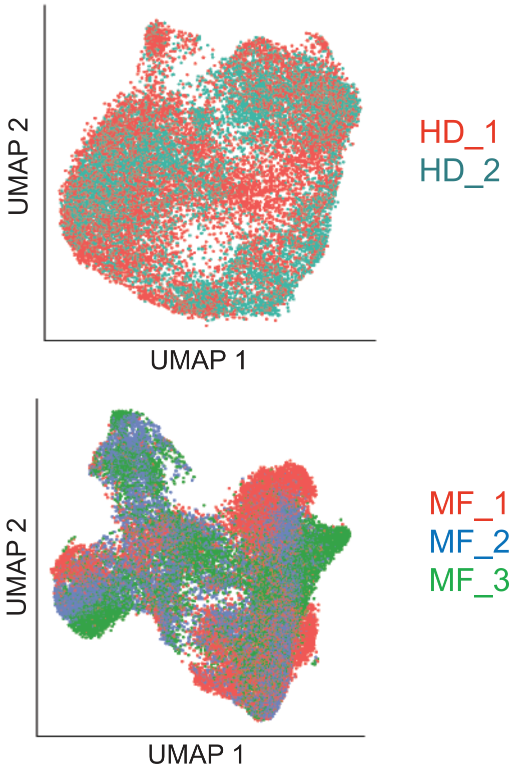


## S1E

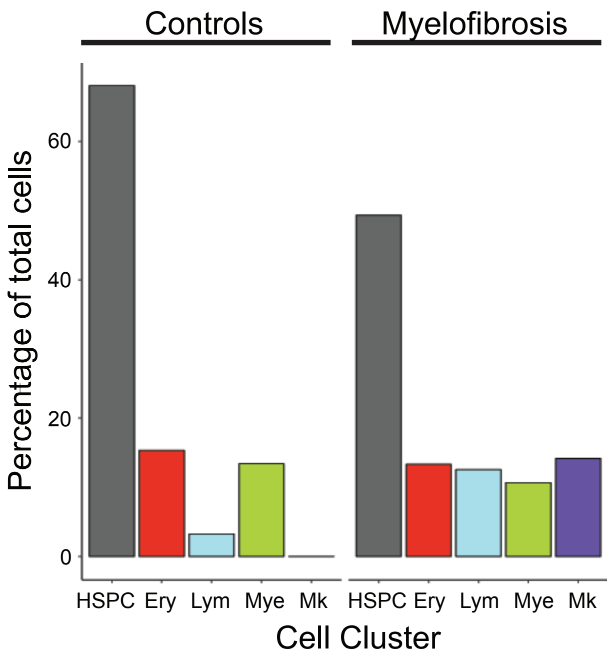

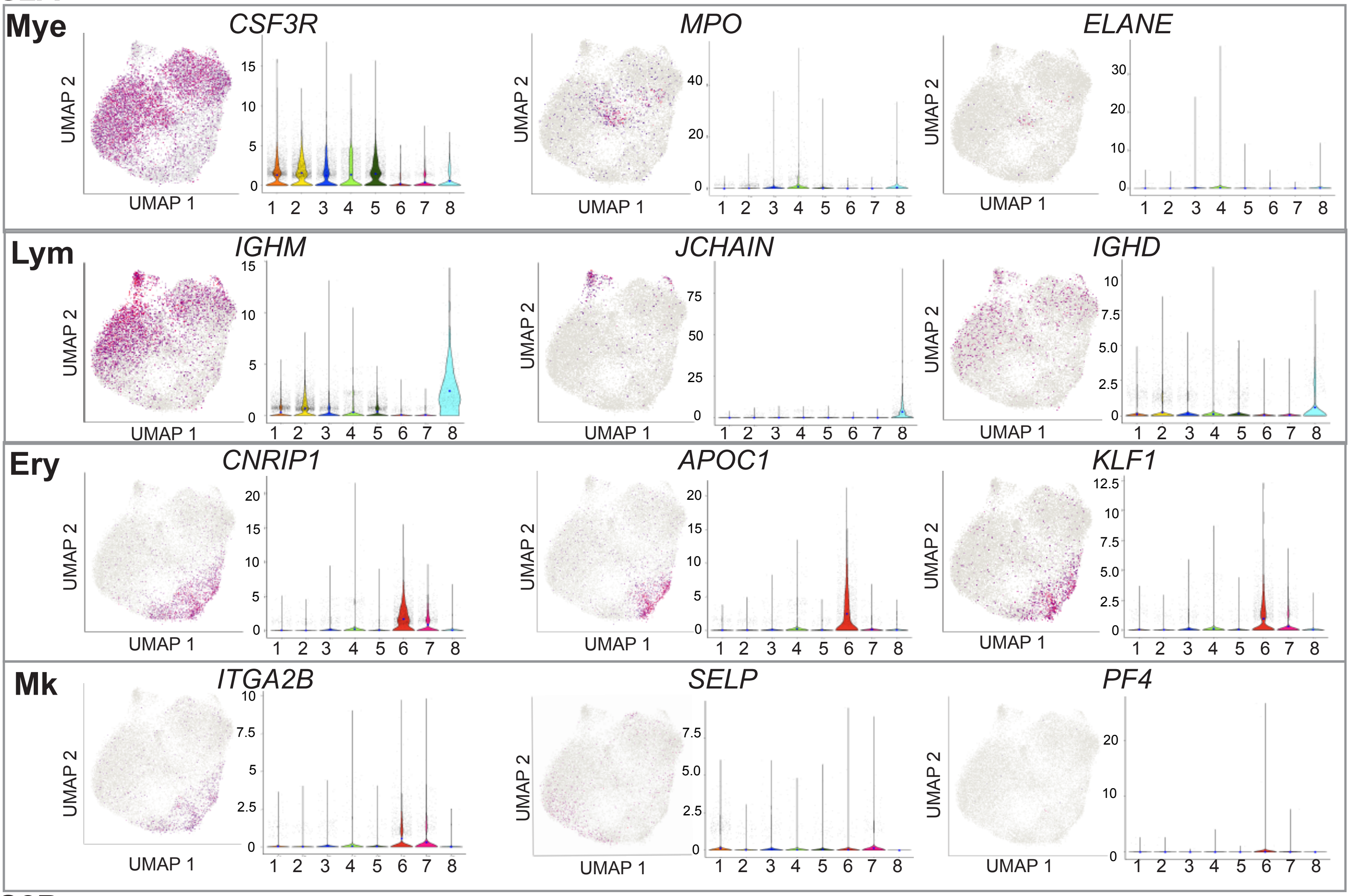

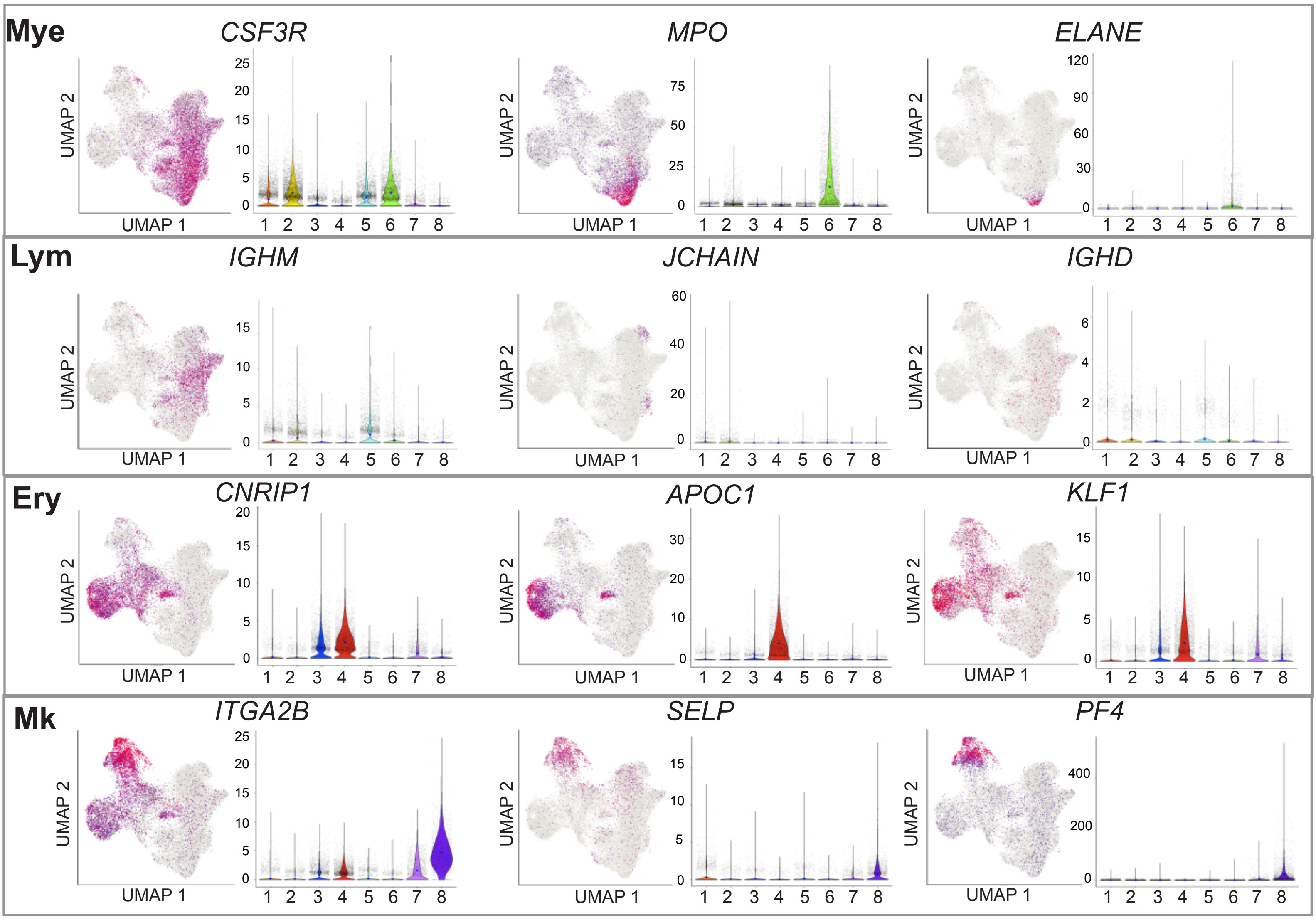

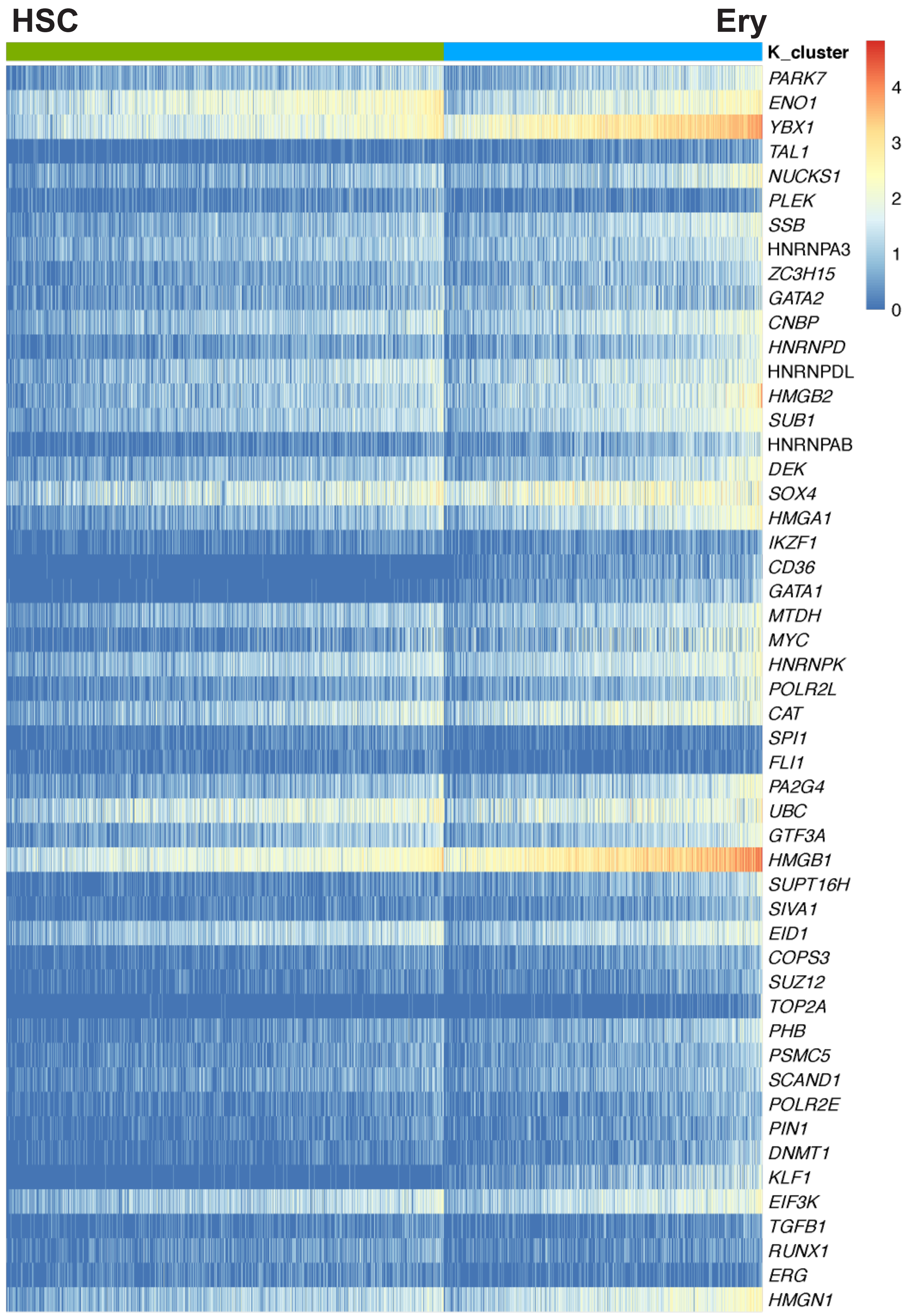

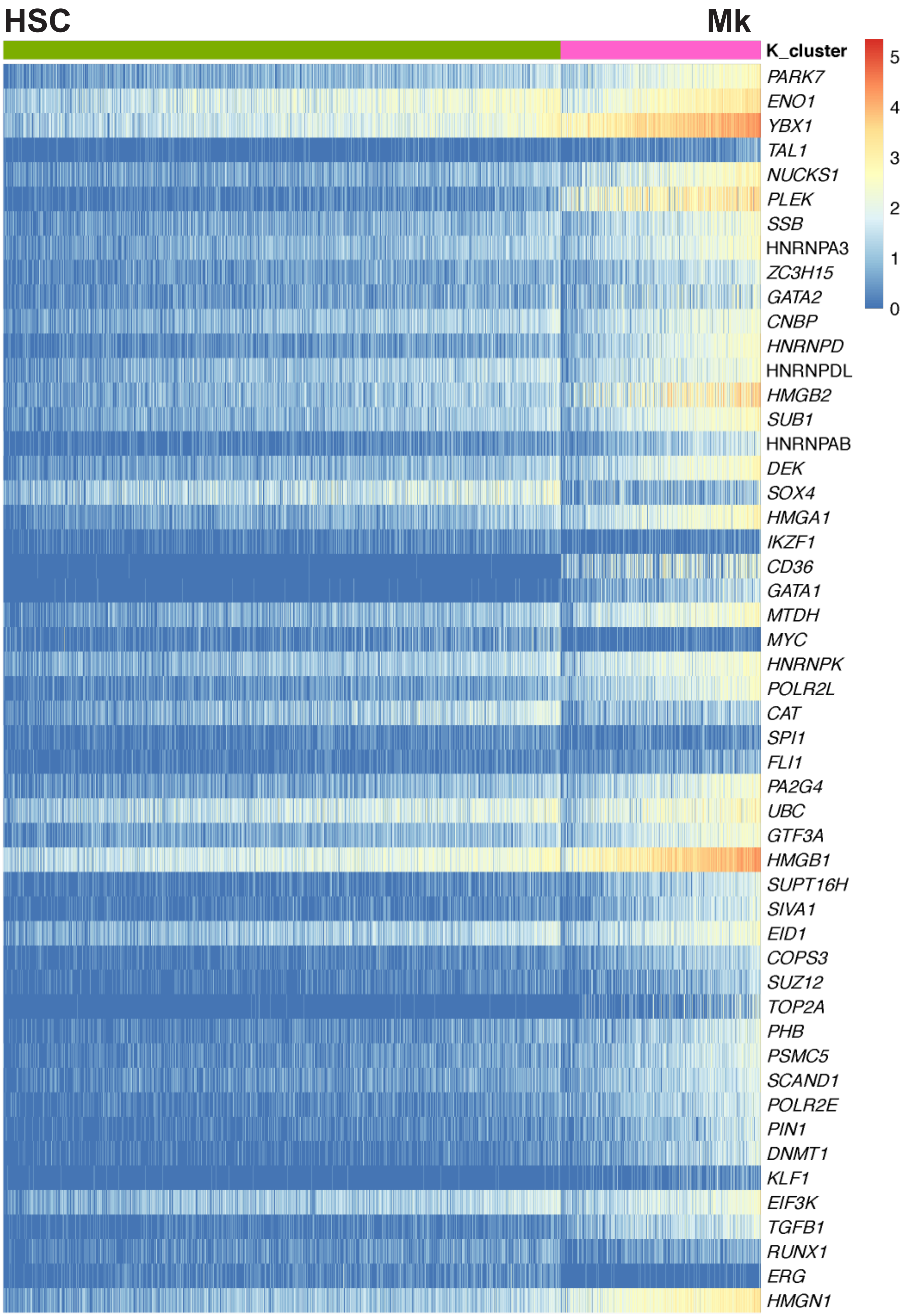

S5A

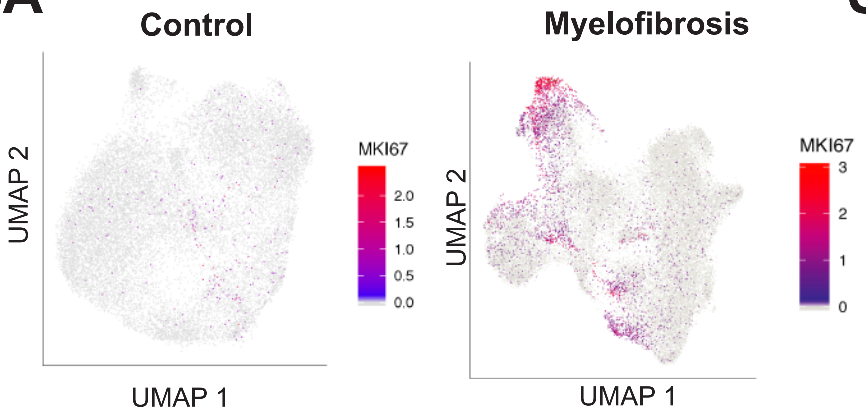


S5B

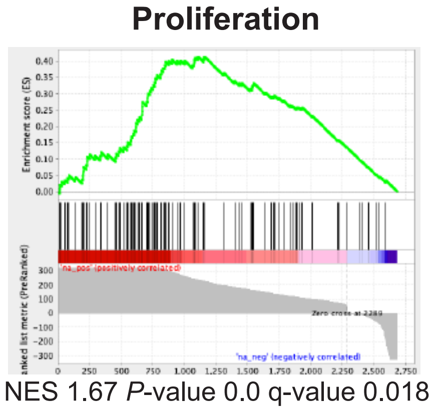


S5C

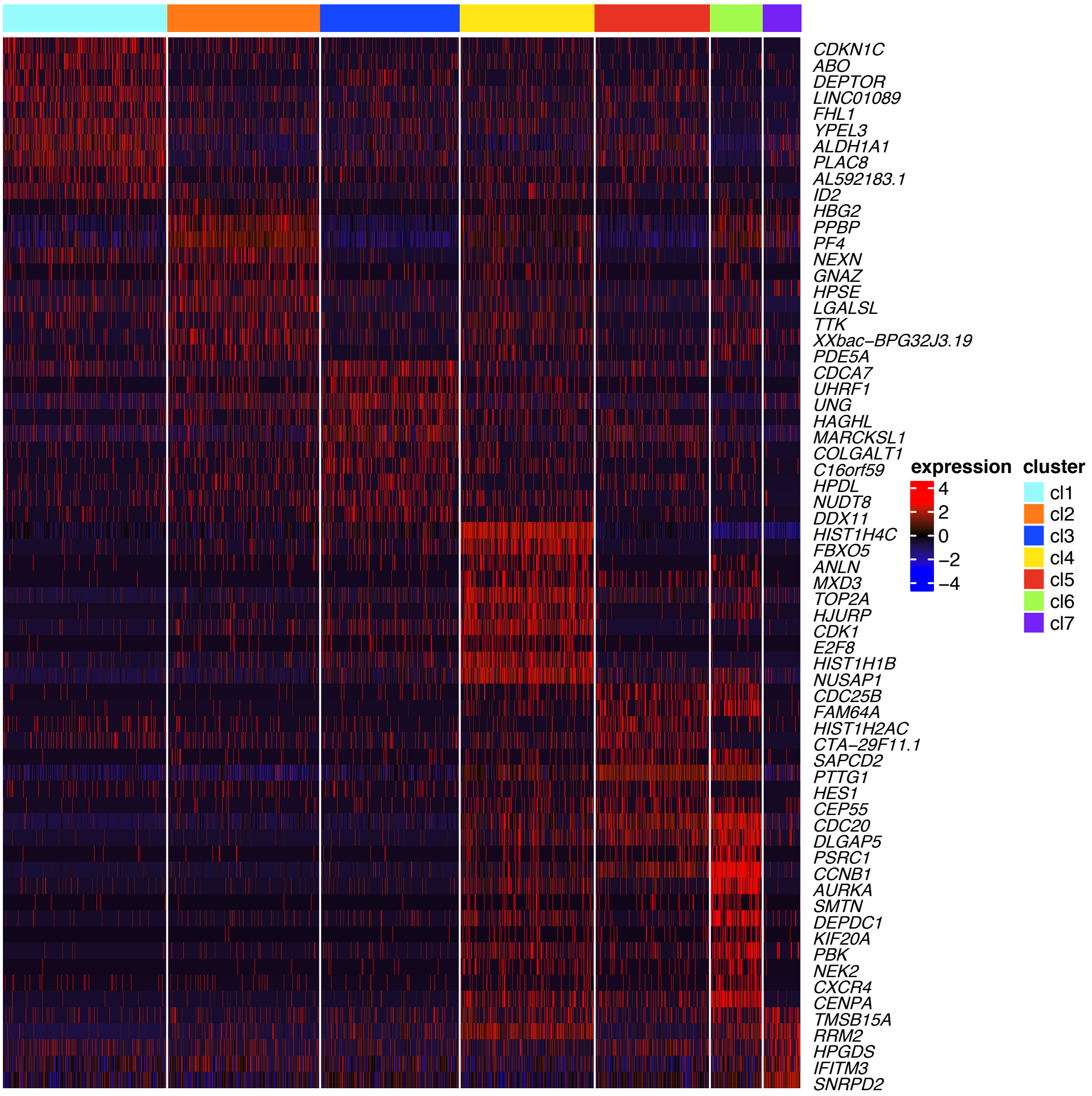


S5D

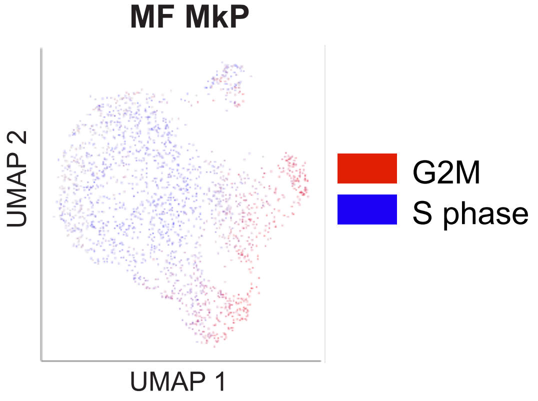

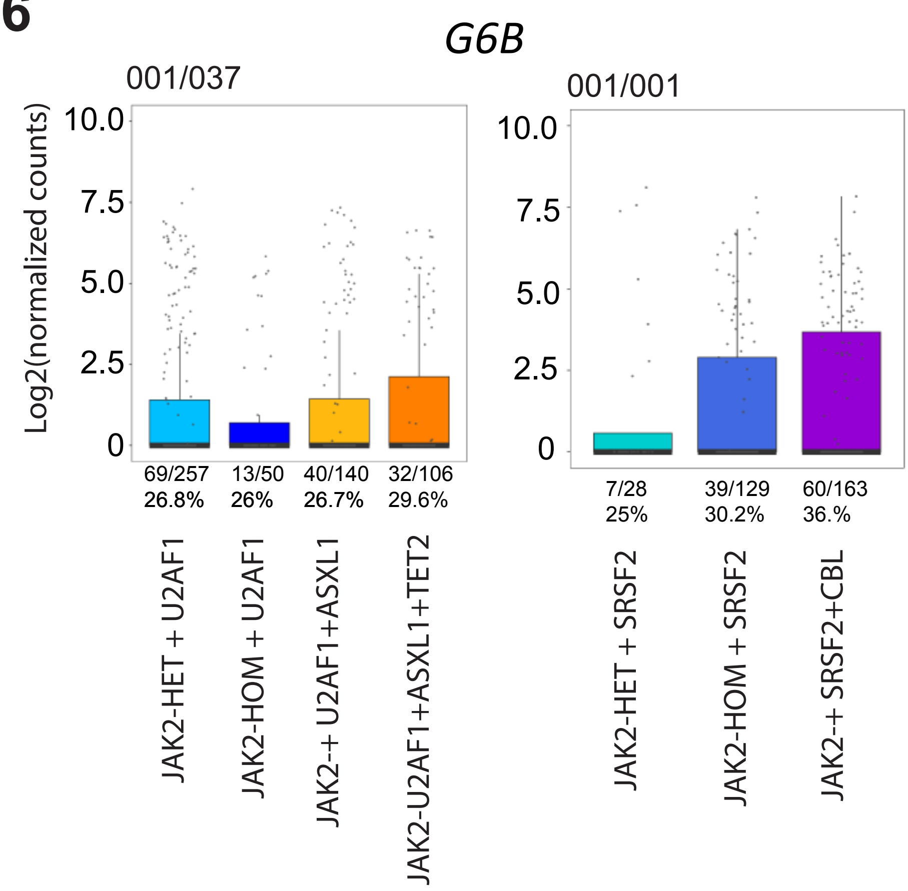

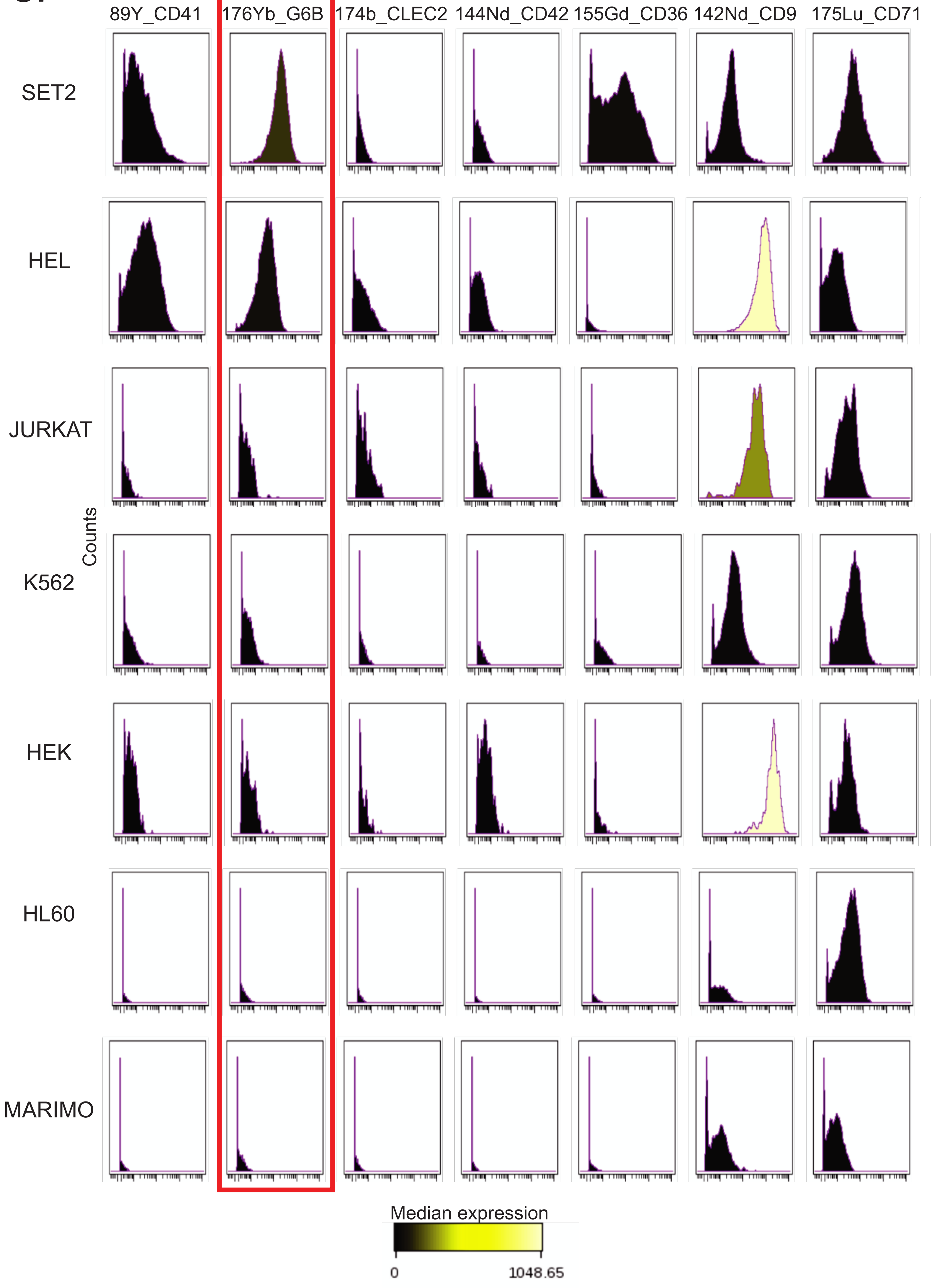
